# Testing for Interactions in Multivariate Data

**DOI:** 10.1101/2025.11.26.690868

**Authors:** Will Penny, Tom Sambrook, Louis Renoult

## Abstract

Factorial designs are a mainstay of the scientific paradigm, allowing the effects of multiple experimental factors and their interactions to be efficiently studied within a single experiment. In brain imaging, however, multivariate data analyses commonly proceed using multivariate decoding and we argue that the standard “difference of accuracies” test is not a true test of interactions. To remedy this situation we propose an encoding method based on a Bayesian Multivariate Linear model which is ideally suited to the factorial analysis of such data. We show how it can be used to test for multivariate main effects and interactions using data from EEG studies of Reward Learning and Declarative Memory. This approach additionally allows for null hypotheses to be accepted and allows one to infer whether multivariate effects are driven by collections of univariate effects or voxel dependencies. We also propose that those wishing to pursue tests for interactions using decoding methods use an “accuracy of differences” test.

**Author summary:** Our understanding of human brain function has been transformed by non-invasive imaging methods such as functional Magnetic Resonance Imaging and Electroencephalography. Statistical modelling has been central to this endeavour with foundational work employing a univariate encoding method in which multiple characteristics of participants behaviour (the stimuli they are exposed to, the decisions they make, the events they remember) are used to predict brain activity at a single point or voxel element (pixel/voxel) in a brain recording, and this process repeats for all voxels in an image. Subsequent work has used multivariate decoding methods which identify categorical behavioural variables from multivariate brain imaging data. Here we propose a new type of multivariate encoding approach in which a Bayesian multivariate linear model is used to predict multivariate image data from multivariate behavioural variables. This approach has three advantages (i) it can correctly test for interactions among experimental factors, (ii) we can quantify the experimental evidence in support of no experimental effects being present and (iii) we can make inferences about the nature of the multivariate effects.

## Introduction

The factorial design is a type of scientific experiment that allows researchers to investigate how multiple factors (independent variables or IVs) influence an outcome (dependent variable or DV). Each factor is tested at distinct levels and the experiment includes combinations of levels across factors. This allows researchers to assess not only how each factor individually affects the outcome, but also how the factors interact with each other. Indeed, an interaction between factors can often be the most crucial finding.

A classic factorial design is the 2-by-2 experiment denoting that there are two factors, each with 2 levels. For example, one factor could be drug intervention (with levels A and B), another factor could be age with levels Young (Y) and Old (O), and the outcome might be health-score. Figure 1 depicts four hypothetical patterns of results. These are extremes presented for purposes of discussion and empirical data will present some mixture of these patterns. Panel D is most relevant for this paper as it gives an example of an interaction without main effects; if the experimenter had not included age as a factor, they would’ve erroneously concluded there was no effect of drug (effects averaged over age are the same for A and B); whereas in fact there is a strong effect - its just signed differently for young versus old participants (producing the so-called crossover interaction).

**Fig 1.**
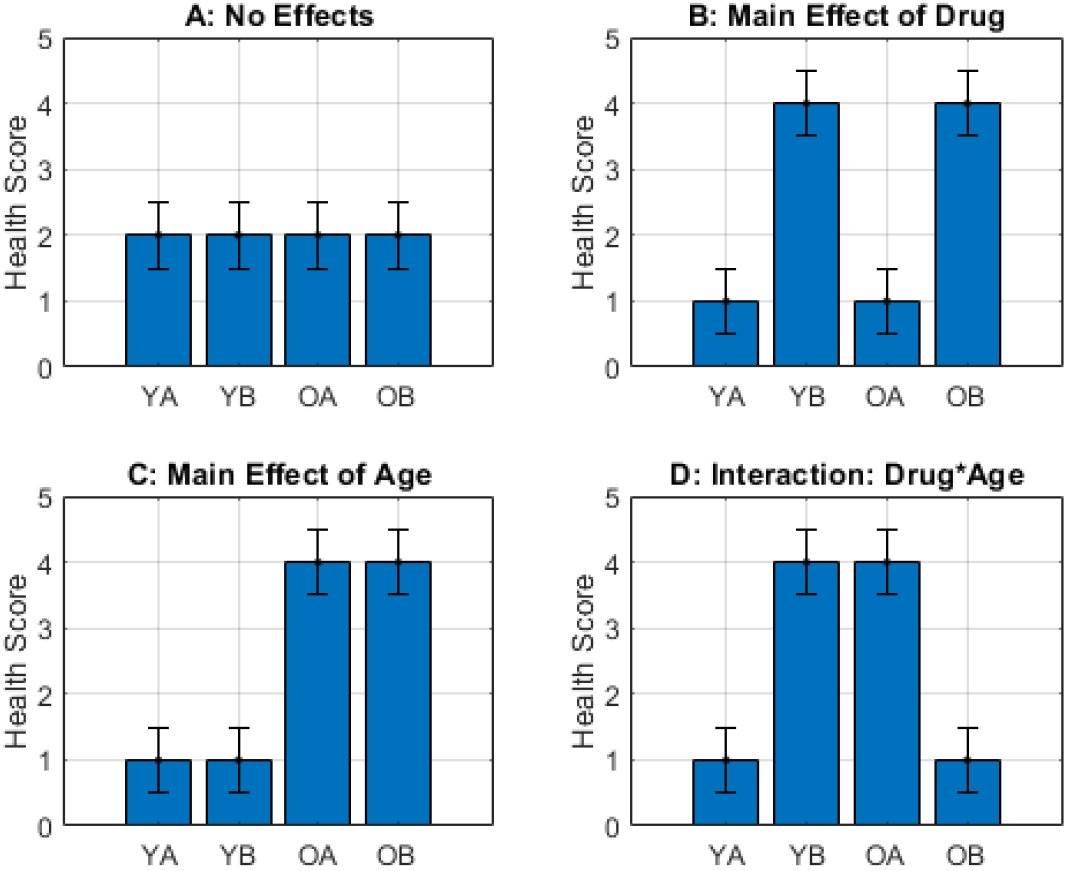
Hypothetical results from a 2-by-2 design. **A:** No significant main effects or interactions, **B:** A main effect of drug *only* - drug B improves the health of both young and old participants relative to drug A, **C:** A main effect of age *only* - both drugs are more effective in older relative to younger participants **D:** An interaction between drug and age *only* - for younger participants drug B is more effective than A, whereas for older participants drug A is more effective than B. By *only* we mean that the other effects are not present.

So far we have considered a single (univariate) experimental outcome - health score. But the same considerations apply when the outcome is multivariate. In the Simulated Data section of this paper we describe the generation of synthetic multivariate EEG data that are analogous to the univariate data depicted in Fig 1. Specifically, Fig 2 is the multivariate equivalent of Fig 1B, and Fig 3 is the multivariate equivalent of Fig 1D.

**Fig 2.**
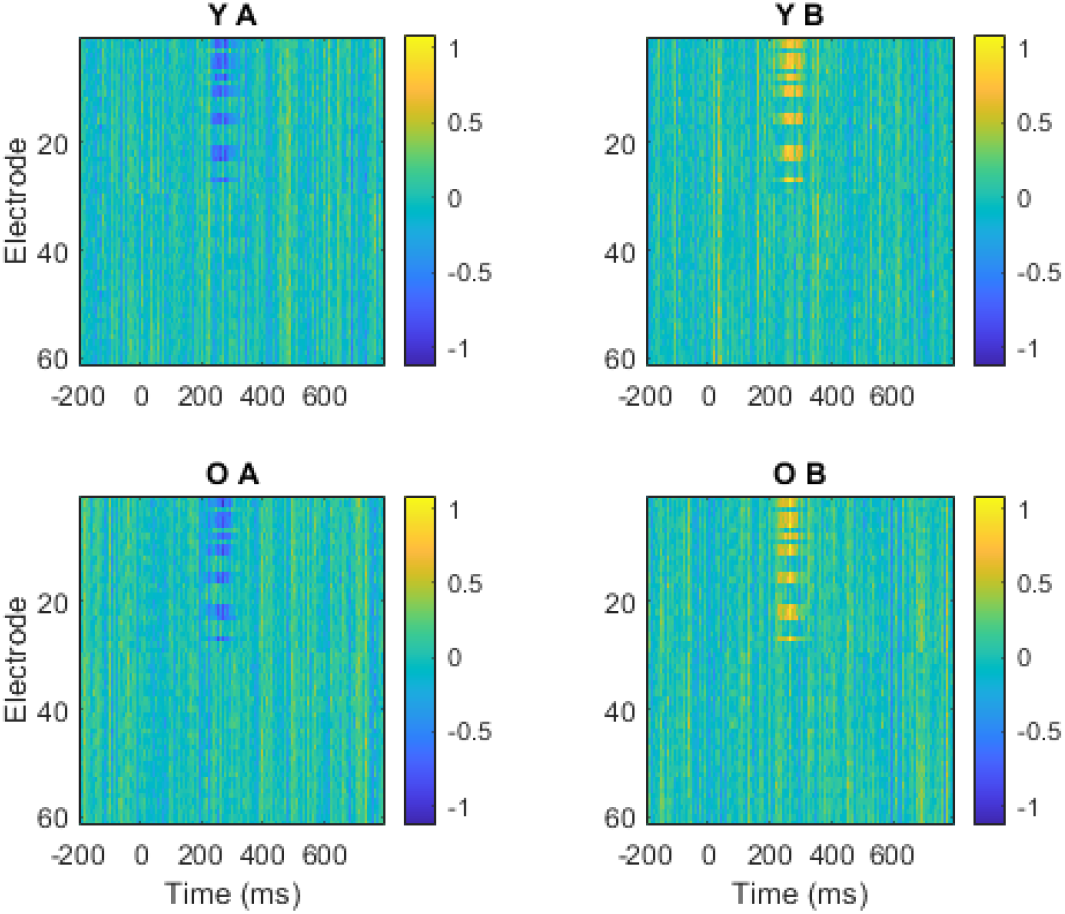
Simulated Data S1. The figure shows the mean responses in the four different conditions with data from Young (Y) participants in the top row, Old (O) participants in the bottom, Drug A in the left column and Drug B on the right. This data set contains an effect of Drug (A versus B) that is manifest in a consistent subset of EEG electrodes centred around 267ms post-stimulus. There is no effect of Group (Old versus Young) or interaction between Drug and Group. This is the multivariate analogue of Fig 1B. This is a high-dimensional data space with data from 61 electrodes and 500 time points. The vertical lines in the images arise from the spatial correlation among electrodes (see Simulated Data section).

**Fig 3.**
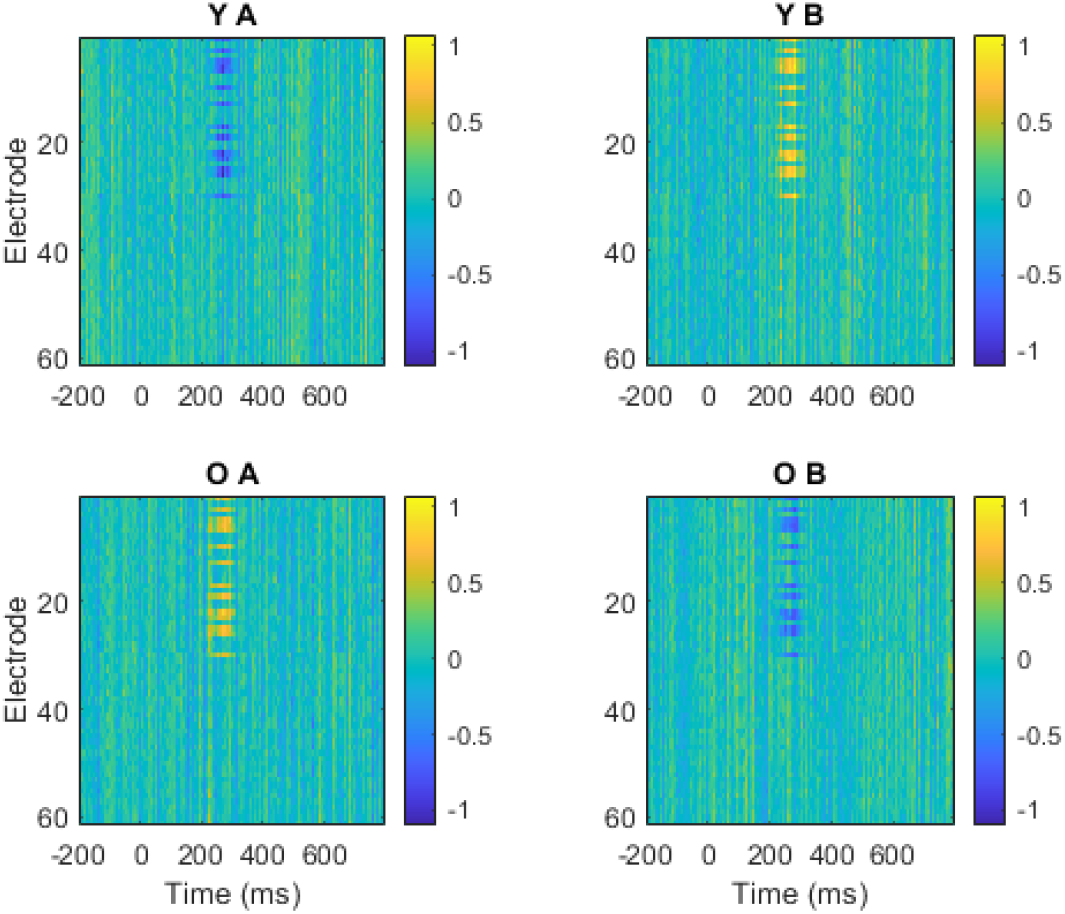
Simulated Data S2. This data set contains an Interaction between Stimulus and Group that is manifest in a consistent subset of electrodes centered around 267ms post-stimulus. There is no effect of Group (Old versus Young) or Drug (A versus B). This is the multivariate analogue of Fig 1D.

**Fig 4.**
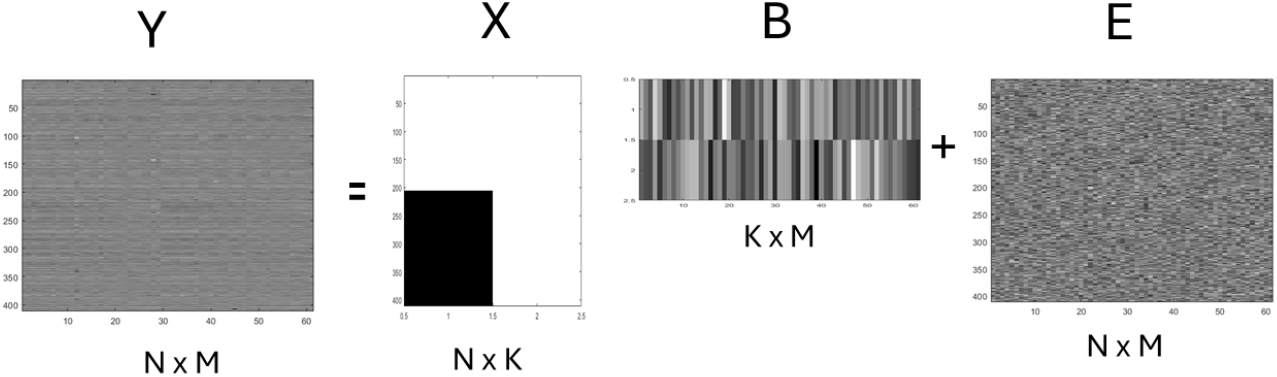
Multivariate Linear Model. A Multivariate Linear Model for making inferences about spatial patterns in EEG data where *Y* are the data, *X* is the design matrix, *B* are regression coefficients, *E* are errors, *N* is the number of trials, *M* is the number of electrodes, and *K* is the number of regression coefficients at each voxel/electrode. This reduces to a Univariate Linear Model if *M* = 1. Here *N* = 400 trials, *M* = 61 electrodes and *K* = 2 regression coefficients. The first column in the design matrix *X* encodes a main effect (difference between first and last 200 trials) and the second column is a vector of ones. The associated regression coefficient vectors encode the size of the main effect (first row) and the average over experimental conditions (second row) at the *M* electrodes. When applied to EEG time series a separate model is used at each time point.

The methodology we present in this paper is able to make the correct inference in each of these cases as shown in Figures 5 and 6, and discussed in more detail in the Results section. Additionally, Bayes factors can be used to correctly accept the null hypothesis which cannot be achieved with Null-Hypothesis Significance Testing (NHST) as this cannot differentiate between a true null and an underpowered study.

**Fig 5.**
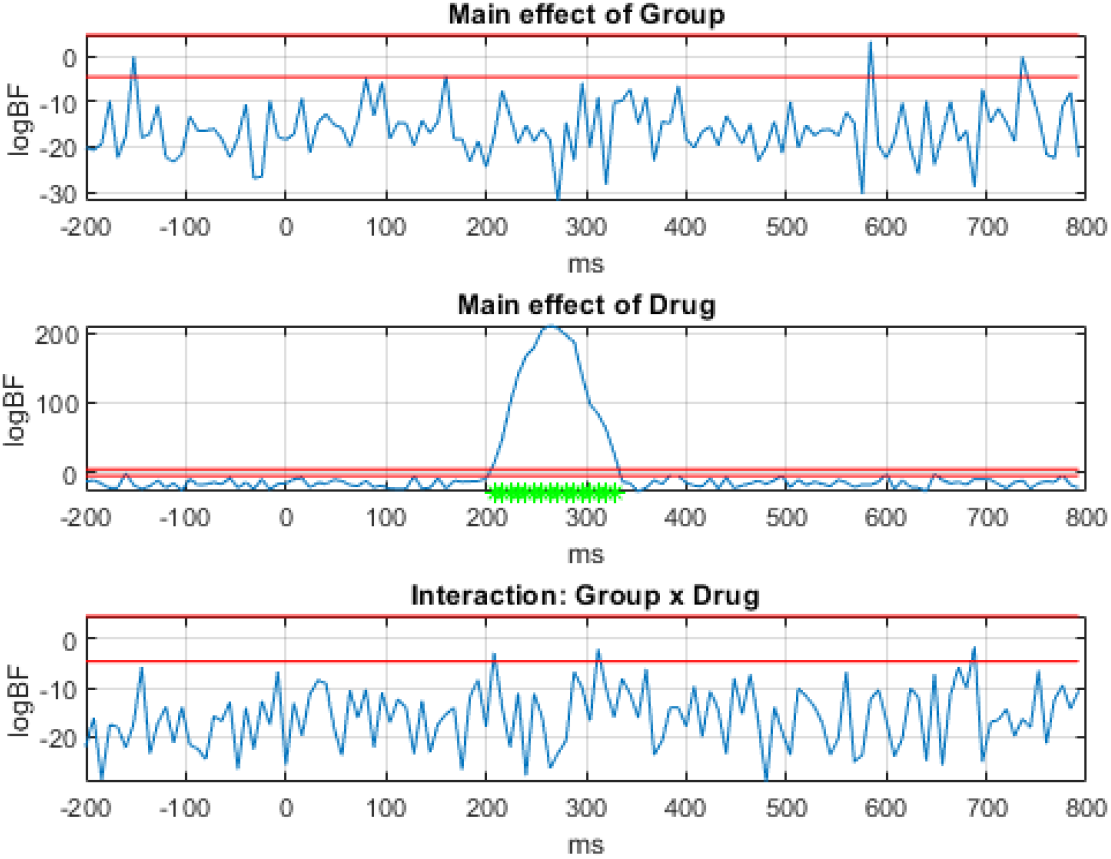
Factorial Analysis of Simulated Data S1. The top to bottom panels plot the Log Bayes Factor in favour of the alternative versus null hypothesis for the main effect of Group (old versus young), main effect of Drug (A versus B) and the interaction. The horizontal red lines are at logBF = −4.6 and logBF=4.6. The alternative hypothesis can be accepted above the top red line, and the null hypothesis below the bottom red line.

**Fig 6.**
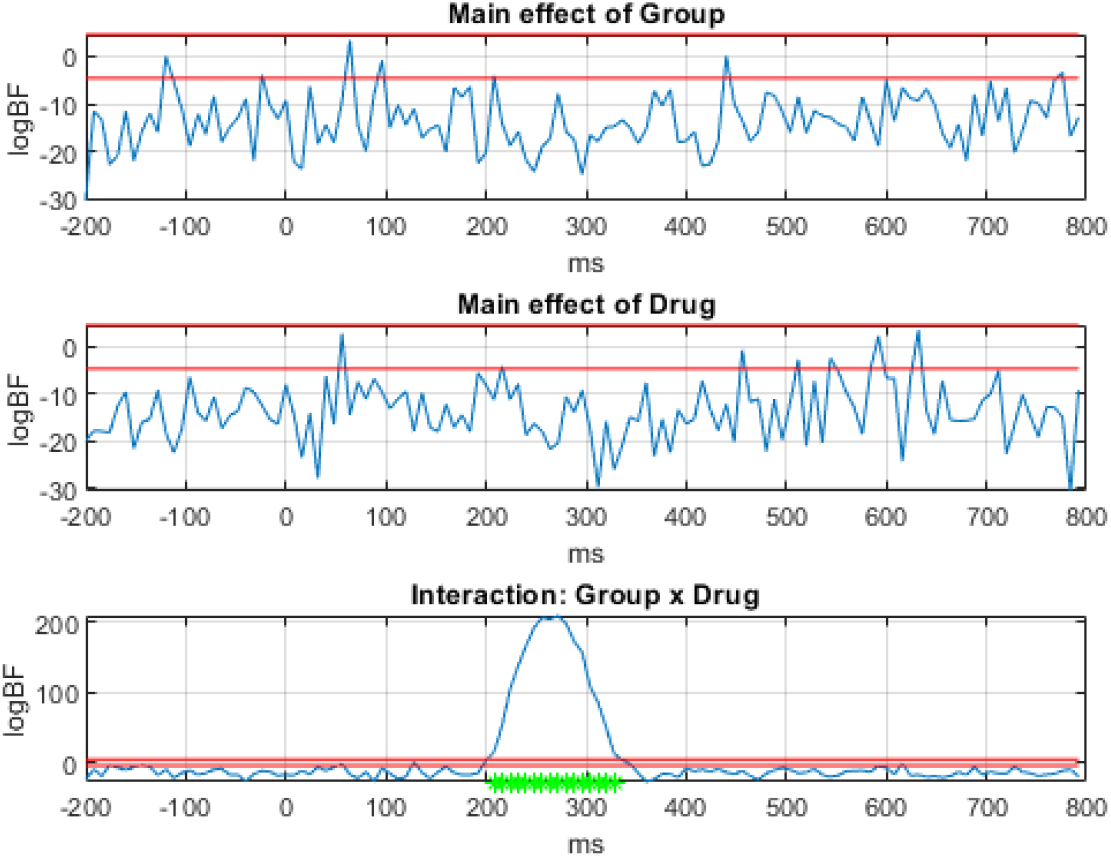
Factorial Analysis of Simulated Data S2. The top to bottom panels plot the Log Bayes Factor in favour of the alternative versus null hypothesis for the main effect of Group (old versus young), main effect of Drug (A versus B) and the interaction. The red lines mean the same as in Fig 5.

In what follows we briefly review the relevant literature on univariate, multivariate, encoding and decoding approaches in neuroimaging. We discuss the application of these methods to EEG and MEG data, and their event-related averages referred to as Event-Related Potentials (ERPs) and Fields (ERFs). Where appropriate we also review relevant work applied to fMRI.

### Mass-Univariate Analyses

ERPs and ERFs comprise data at multiple sensor-by-time locations. Each location is also referred to as a “space-time point” or voxel (volume element). In mass-univariate analyses, experimental effects are initially assessed separately at each voxel but the resulting statistical inference must then account for the multiple statistical comparisons across all voxels. Whilst a Bonferroni correction is possible (multiplying p-values by the number of comparisons) the resulting inferences are underpowered. This is because data at neighbouring space-time points are highly correlated, and not independent as assumed by Bonferroni. Better corrections can therefore be made by taking this correlation into account, and this can be implemented parameterically using Random Field Theory (RFT) [1, 2], or non-parametrically using Nonparametric Cluster Inference (NCI) [3].

Whilst both of these tools were motivated by addressing the multiple comparisons problem for univariate models, they both end up viewing the data as a multivariate quantity. For example, significant clusters found by NCI are just that - it’s the clusters of locations that are significant, not specific voxels within them. Similarly, the set- and cluster-levels of inference in RFT both pertain to distributed patterns of responses. Chumbley et al [4] take this further by proposing that RFT be used to test whether the local density of responses is higher in one brain region than another, thus allowing inferences about functional segregation (region 1 is involved in task X but region 2 is not). This approach has been applied to fMRI data.

### Multivariate Decoding

Generically, decoding methods predict behaviour from brain activity whereas encoding methods predict brain activity from behaviour. The dominant approach to Multivariate Analyses of M/EEG data is to use a decoding approach, otherwise known as Multivariate Pattern Analysis (MVPA), and to consider the signal to be multivariate over space, but univariate over time. These analyses generally provide plots of classification accuracy as a function of peri-stimulus time and are supported by tutorials [5] and freely available libraries of software tools [6–8].

Most often, experimental conditions are decoded using information from all MEG sensors or EEG electrodes. For example, Cichy and Oliva used Support Vector Machines (SVMs) for MEG decoding to identify the time course of human object recognition [9]. In other applications a subset of (pre-specified) electrodes are used. For example, Mares et al. investigate face processing in adults and children [10] using patterns of activity over occipito-temporal electrodes.

It is also possible to identify which sensors contribute significantly to decoding. For example, Kurth-Nelson et al. [11] used an SVM approach to identify time points at which patterns could be recalled in an MEG-based associative retrieval task. Here, maps of the contribution of each sensor to the overall classification performance were computed by randomly sub-sampling sensor space and reporting the average accuracy for samples that included that sensor. However, Haufe et al. [12] provide compelling examples and mathematical analysis to show that the weight vectors used in decoding algorithms, such as Linear Discriminant Analysis (LDA), cannot be interpreted to mean that electrodes with significantly non-zero weights contain (univariately) decodable information.

King et al. [13] review what they refer to as the “temporal generalisation” method in which classifiers are trained on one time point but tested on all other time points. This is also known as “temporal cross-decoding” [14, 15] and allows for the identification of processes that are either common or differentiated across time thereby helping to unpack how neural representations are manipulated and transformed.

The above approaches all classify time points independently. This is an optimal approach if the main goal of the analysis is to find out at which time points various mental representations are active. This makes sense as the main strength of EEG/MEG as compared to fMRI is to identify *when* various neural or cognitive processes occur.

However, if the overall goal is (a) to better understand the ERP signal including potential dependencies between components or (b) to assign participants to clinical groups for ERP-based clinical diagnostics, then it is better to combine information from multiple time points. One example of this approach is taken in Vahid et al. [16] in which a deep neural net learnt to use information from multiple time points in single-trial EEG epochs - principally the N1 (190-250ms), N2 and lateralised readiness potential (LRP) (both 300-400ms) - to accurately categorise experimental conditions in an action control task.

Friston et al. [17] have developed Multivariate Bayesian (MVB) decoding models for the analysis of multivariate fMRI data. These provide a mapping from features of multivariate spatial patterns (e.g. distributed or clustered patterns) expressed in fMRI time series in local brain regions, via hemodynamic deconvolution, to experimental condition labels. MVB has been used, for example [18], to show that neural responses during episodic memory tasks are more spatially distributed in older versus younger participants.

### Interactions

The standard way to test for interactions using multivariate decoding is what we shall refer to as a “difference of accuracies” test. This proceeds by testing whether the decoding accuracy, between the levels of one factor (such as Drug), changes as a function of the level of another factor (such as Group). This test will fail on the interaction effects in the S2 data (see Fig 3) because the decoding accuracies are not different in the young versus old groups. Here, the only differences are in the sign of the effects and not the magnitudes. This may perhaps seem like an esoteric case but differences in signs of ERP signals are readily produced by small changes in the location (and therefore orientation) of underlying equivalent dipoles. Additional scenarios causing the “difference of accuracies” test to fail are when the second factor simply rotates the decision boundary. This can have substantial effects on the nature of the underlying signals but no effect on discrimination accuracy.

However, the “differences of accuracies” test will not always fail and there are very many completely valid empirical findings of interactions revealed by this method. For example, Smith et al. [19] examined the effect of task (with two levels - explicit or incidental) on the decoding of facial identity (with six levels corresponding to 6 different individuals). They found a significant reduction in decoding accuracy, at specific peri-stimulus time points, when the task was incidental rather than explicit. This must have been driven by underlying differences in the multivariate patterns - that is, an interaction.

Smith et al. [19] also tested for an interaction between task and facial expression (with 7 levels - happy, sad, tearful, disgusted, angry, surprised and neutral) but found no differences in decoding accuracy as a function of task. But as we have argued with the simulated data there can still be differences in multivariate patterns even when there are no differences in decoding accuracy. Overall, therefore, “difference of accuracies” is not a satisfactory test for interactions because negative results are uninterpretable.

The “difference of accuracies” approach has also been adopted in methods that use Representational Similarity Analysis of EEG and MEG data as shown for example in Dobbs et al. [20], Ambrus et al. [21] and Xu et al [22]. The overall approach here is to first create an empirical Representational Dissimilarity Matrix (RDM) where each entry in the matrix is cross-validated decoding accuracy between multivariate EEG/MEG responses to a pair of stimuli (each stimulus being shown many times) and then to regress this (vectorised) RDM onto several model RDMs each reflecting the dissimilarities that are expected to arise under each factor (e.g. for the factor of gender, dissimilarity would be 1 for pairs of faces from different genders and 0 for the same gender). Further model RDMs are included as independent variables so their effects can be partialled out. The partial correlation between the empirical RDM and the model RDM is then the measure of accuracy. Tests are then made as to whether these accuracies are different between levels of a second factor. This is therefore also a “difference of accuracies” test.

We return to this issue in the Discussion where we propose an alternative “accuracy of differences” method for testing for interactions using multivariate decoding. In this paper, however, we pursue an alternative method based on a Multivariate Linear Model (see below).

### Encoding Models

Whereas multivariate decoding models predict category labels from imaging data, encoding models predict imaging data from category labels (and potentially other variables). Encoding models have been developed for analysing patterns of fMRI activity from visual cortex recorded whilst participants made decisions about the orientation of visual gratings [29]. This rested on the specification of an encoding model that specified orientation tuning in a population of neurons, with additional weight vectors that mapped these responses onto voxel activations in the brain region of interest. Model parameters were estimated from empirical data which then allowed for a read-out of a participant’s decision uncertainty. Similar encoding models, that also make use of hypothesised population responses to relevant stimulus features, have been proposed for analysing patterns in EEG data. This topic is reviewed in Holdgraf et al [30] with an emphasis on linear encoding models and auditory tasks.

Building on previous work that views EEG signals as arising from the dynamical evolution of a small number of discrete neural states, Higgins et al. [31] propose a spatio-temporal encoding model of EEG data. This uses a left-right hidden Markov model for the temporal evolution of the states, and each state has spatial parameters that project to EEG electrodes. Model parameters are allowed to vary with experimental condition (encoding) and Bayes rule is then used to infer the experimental condition from novel EEG data (decoding). The Variational Bayes framework [32] is used for model estimation and inference and the overall algorithm is shown to provide better decoding performance than standard approaches. The main strength of this approach is that the timing of the state activations can vary over trials, a feature which was shown to be well-suited to empirical data.

### Multivariate Analysis of Variance

The standard statistical approach for testing whether multivariate patterns differ over experimental conditions is Multivariate Analysis of Variance (MANOVA). Inference under parametric assumptions is based, for example, on the use of Pillai’s trace statistic and its approximate F-distribution [23]. MANOVAs have been previously used in ERP research, for example, to test for group differences in the pattern of activity over a small number of midline electrodes during the P300 elicited in a visual oddball paradigm [24].

Allefeld and Haynes have applied MANOVA to pattern analysis of fMRI data [25]. Rather than relying on parametric results for statistical inference, they instead use a performance metric called pattern distinctiveness, D, and use cross-validation to provide an unbiased estimate of D from empirical data. They argue that D is to be preferred to classification accuracy as a performance metric as it can be applied in two important novel contexts; (i) to factorial experimental designs thereby allowing for inferences about interactions and (ii) to multivariate models with design matrices containing continuous as well as discrete covariates, and point out that there is no classification accuracy equivalent in either of these contexts. This important point partly motivated the modelling work in this paper. In related work, the relative merits of encoding versus decoding approaches are reviewed at length in [26] with a focus on the nature of the underlying brain responses that are driving experimental effects.

Friston et al. [27] describe a multivariate analysis of MEG data comprising two stages. First, spatio-temporal data are subject to a singular value decomposition. Second, the resulting spatiotemporal modes are subject to a MANCOVA (MANOVA with covariates [28]). This approach was applied to single-trial MEG data and identified pre-stimulus effects in a motor learning task.

### The Present Study

In this paper we propose that inferences about main effects and interactions in multivariate imaging data be made using a Bayesian Multivariate Linear model with Conjugate priors which we refer to as MLC. Figure 4 shows how MLC is configured to make inferences about spatial patterns over EEG electrodes. The corresponding Bayesian inference procedures are described in the Methods section below. We first test this approach using Simulated Data Sets S1, S2 and S3. Data sets S1 and S2 contain homogeneous effects (same effects at each sensor) so as to make a clear comparison with the univariate data discussed earlier in the introduction. Data set S3 contains heterogeneous effects. We then test the approach using a within-subject analysis of Reward Learning ERPs, and a between-subject analysis of Declarative Memory ERPs. We compare the inferences made with those provided using decoding methods based on Support Vector Machines, Linear Classifiers and k-nearest neighbour classifiers (but only for the make effects because, as argued above, we regard negative-findings about interactions to be uninterpretable). The MLC approach has three characteristics that may make it especially useful to neuroimaging researchers (i) it provides a valid test for interactions, (ii) it can be used to accept the null and (iii) it can be used to identify the underlying nature of the multivariate effects, that is, whether or not inferences are driven by a collection of univariate effects or voxel dependencies. These characteristics and full details of the data sets and provided in the Materials and Methods section below.

## Results

### Simulated Data

We fitted simulated datasets S1, S2 and S3 using the Multivariate Linear Model with Conjugate Priors (MLC). Fig 5 plots the resulting inferences for S1 with the alternative hypothesis correctly accepted for the main effect of Drug at *t* ∼ 267ms, and the null hypotheses correctly accepted for the main effect Group and the interaction. Fig 6 plots the resulting inferences for S2 with the alternative hypothesis correctly accepted for the Interaction at *t* ∼ 267ms, and the null hypotheses correctly accepted for the main effects Group and Drug. Fig 7 plots the resulting inferences for S3 with the alternative hypothesis correctly accepted for the main effect of Drug at *t* ∼ 267ms and the interaction between Group and Drug at *t* ∼ 533ms. The null hypotheses are correctly accepted at other time points. There are several time points in each of the above figures with `LogBF` between the red lines, indicating that neither the null or alternative hypotheses can be accepted.

**Fig 7.**
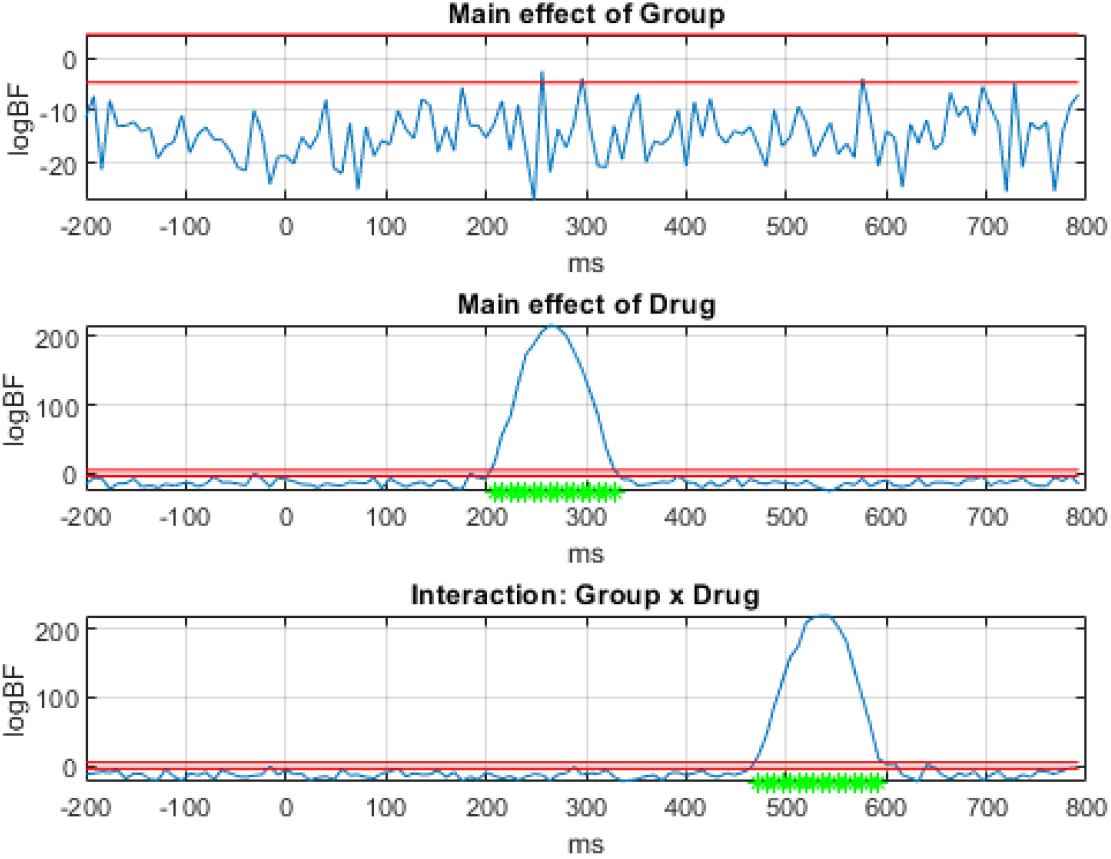
Factorial Analysis of Simulated Data S3. The top to bottom panels plot the Log Bayes Factor in favour of the alternative versus null hypothesis for the main effect of Group (old versus young), main effect of Drug (A versus B) and the interaction. The red lines mean the same as in Fig 5.

Table 1 shows the total computation time on a desktop PC (Hewlett Packard Z2 Tower G9 workstation with 16Gb RAM) on data sets generated using the procedure as for S1, but additionally varying the number of electrodes, *M*. We note that MLC is about five times faster than SVM for data with 64 electrodes but about 20 percent slower with 320 electrodes. We return to this issue in the Discussion.

**Table 1.** Total Compute Time (seconds). For the Multivariate Linear model with Conjugate priors (MLC) and the Support Vector Machine (SVM) with 10-fold cross-validation as we vary the number of electrodes.

| Number of Electrodes | MLC | SVM |
| --- | --- | --- |
| 64 | 4.9 | 21.4 |
| 128 | 18.7 | 23.6 |
| 192 | 30.1 | 36.8 |
| 256 | 50.9 | 51.2 |
| 320 | 77.1 | 63.3 |

### Reward Learning

This section presents a factorial analysis of the Reward Learning data. We use data from a single subject only as we wish to demonstrate how MLC can be used for a within-subject analysis. As before, we implemented multivariate tests at each peristimulus time point. We fitted MLC to the EEG epochs from this subject and tested for the main effect of Reward Prediction Error (RPE) sign (negative versus positive), main effect of RPE magnitude (small versus large) and the interaction between these factors with results shown in Fig 8.

**Fig 8.**
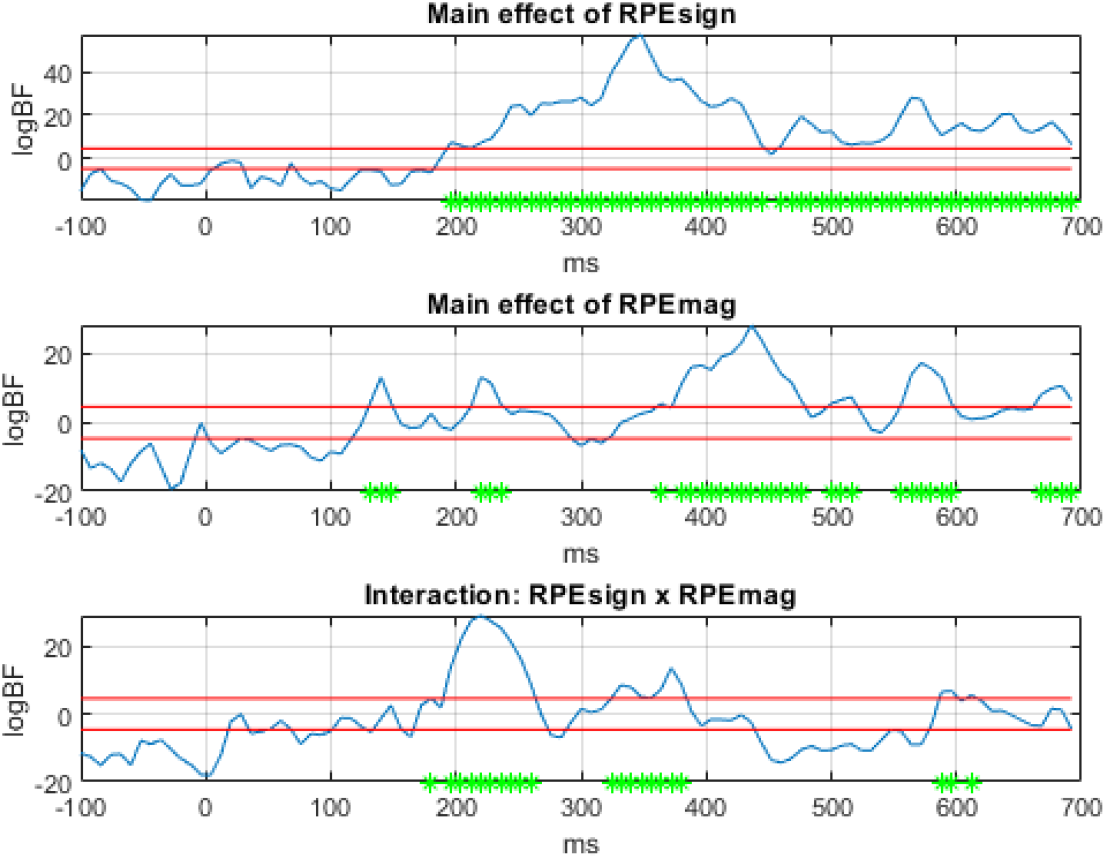
Factorial Analysis of Reward Learning. The top to bottom panels plot the Log Bayes Factor in favour of the alternative versus null hypothesis for the main effect of Reward Prediction Error sign (negative versus positive), main effect of RPE magnitude (small versus large) and the interaction. The red lines mean the same as in Fig 5.

The results show that there are no multivariate effects until approximately 150ms post-stimulus. The log Bayes factors prior to this are strictly negative, indicating that the null hypothesis is to be accepted. From 200ms onwards, the main effect of `RPEsign` is mostly significant with a peak at 348ms and the main effect of `RPEmag` is significant more intermittently with a peak at 436ms. The interaction effect is more localised with a broad peak effect at 220ms. Figure 9 uses voxel-wise ANOVAs to create sensor maps of the univariate effects at the peak times of the corresponding multivariate effects. These maps indicate that there are indeed univariate effects at the times of the strongest multivariate effects.

**Fig 9.**
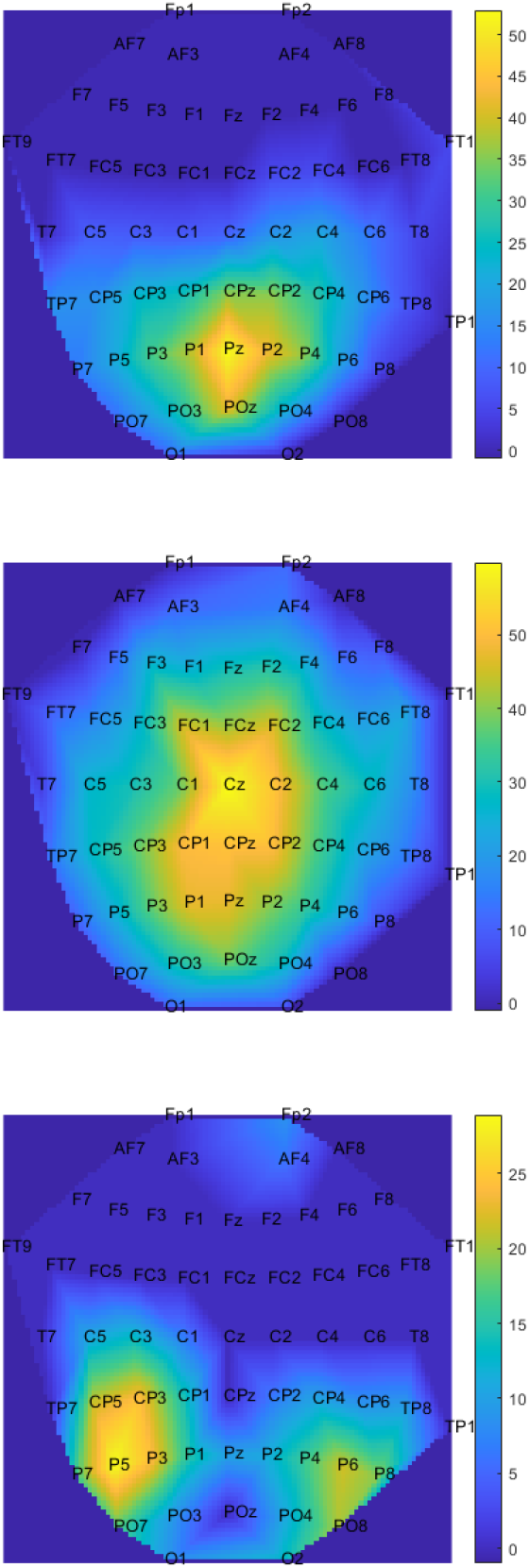
Spatial Maps at Peak Effect Times. The maps show the main effect of `RPEsign` at 348ms (top), the main effect of `RPEmag` at 436ms, and the interaction at 220ms. The colours indicate the value of the F-statistic from a voxel-wise ANOVA. F-values are set to zero for voxels that do not satisfy an FDR correction over electrodes (*FDR <* 0.05).

For comparison with the MLC results, Fig 10 presents a decoding analysis of the main effects using Support Vector Machines (SVMs) which found peak decoding accuracy for `RPEsign` at 348ms and for `RPEmag` at 220ms. MLC and SVM are broadly in agreement. Overall, as shown in Tables 2 and 3, the correlation between MLC Bayes Factors and SVM classification rates was highly significant and the strength of correlation was of a similar magnitude as between different decoding algorithms. For making inferences about main effects, the pattern of findings from MLC is most correlated with SVM and linear decoders, rather than kNN classifiers.

**Table 2.** Correlations among encoding and decoding scores for main effect of RPEsign (negative versus positive). In the top row the correlations are between the log Bayes factor time series from MLC (Fig 8) and the *p*_*corr*_ time series from the decoders (Fig 10 for SVM). All other correlations are among *p*_*corr*_ time series. All correlations are significant at *p <* 0.001.

| Encoder/Decoder | Decoder |  |  |  |
| --- | --- | --- | --- | --- |
|  | kNN | Linear | SVM | PCA-SVM |
| MLC | 0.64 | 0.92 | 0.94 | 0.86 |
| kNN | - | 0.59 | 0.64 | 0.57 |
| Linear | - | - | 0.94 | 0.82 |
| SVM | - | - | - | 0.82 |

**Table 3.**
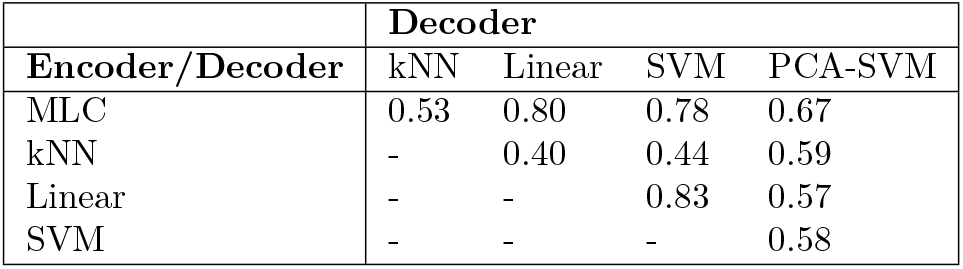
Correlations among encoding and decoding scores for main effect of RPEmag (small versus large). In the top row the correlations are between the log Bayes factor time series from MLC and the *p*_*corr*_ time series from the decoders. All other correlations are among *p*_*corr*_ time series. All correlations are significant at *p <* 0.001.

**Fig 10.**
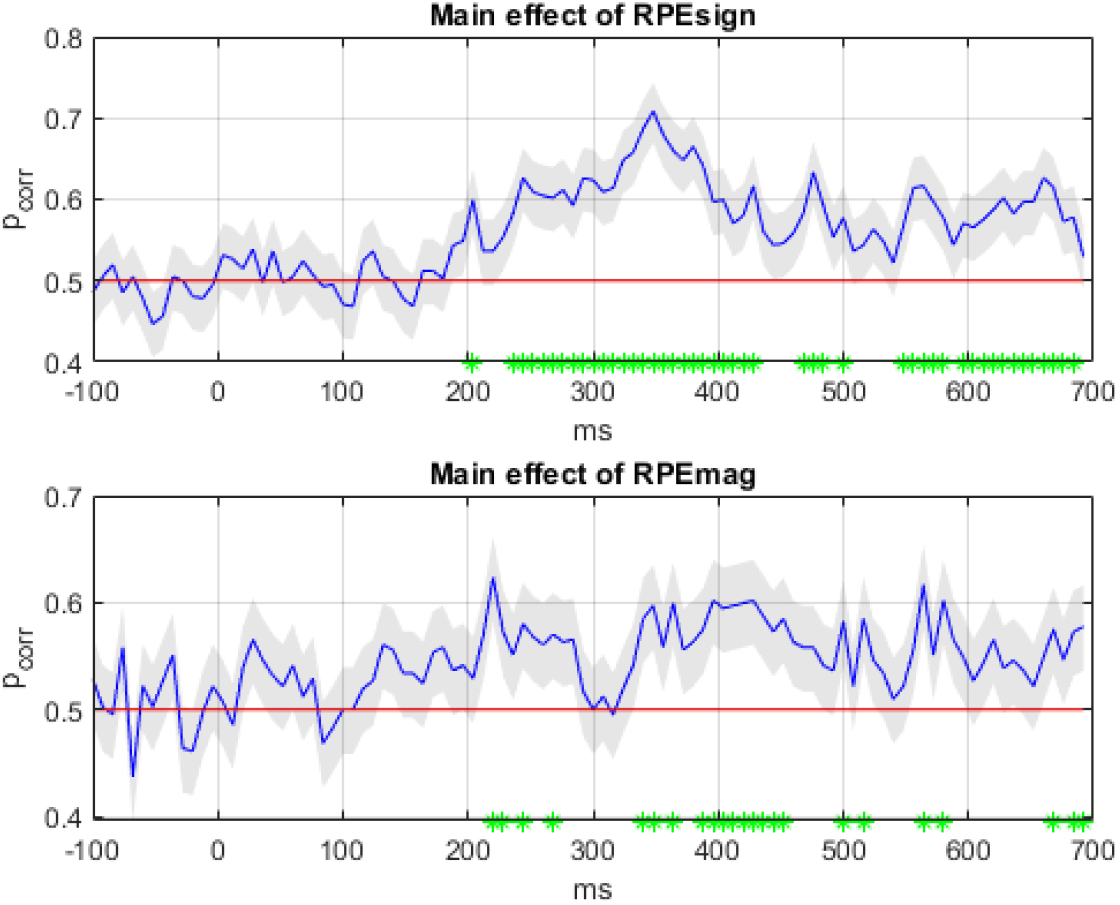
Decoding Analysis of Reward Learning data. The panels plot the decoding accuracy, *p*_*corr*_, with 90 percent confidence intervals for the main effect of Reward Prediction Error sign (negative versus positive) and the main effect of RPE magnitude (small versus large) using an SVM decoder. The green asterisks denote significant decoding periods (*FDR <* 0.01).

### Declarative Memory

This section presents a factorial analysis of Declarative Memory ERPs by implementing multivariate tests at each peristimulus time point. We focus on Memory-Type, defined as a semantic versus episodic distinction. MLC was fitted to data from 26 subjects in each Group (old and young) and we tested for the main effect of Group (old versus young), main effect of Memory-Type (semantic versus episodic) and the interaction between the two, with results shown in Fig 11. The results show that there are no multivariate effects until approximately 100ms post-stimulus. Log Bayes factors prior to this are negative, indicating that the null hypothesis is to be accepted. From this point on, the main effect of Group is significant, and the main effect of Memory-Type within two clusters of time points. The interaction is significant from about 570ms onwards.

**Fig 11.**
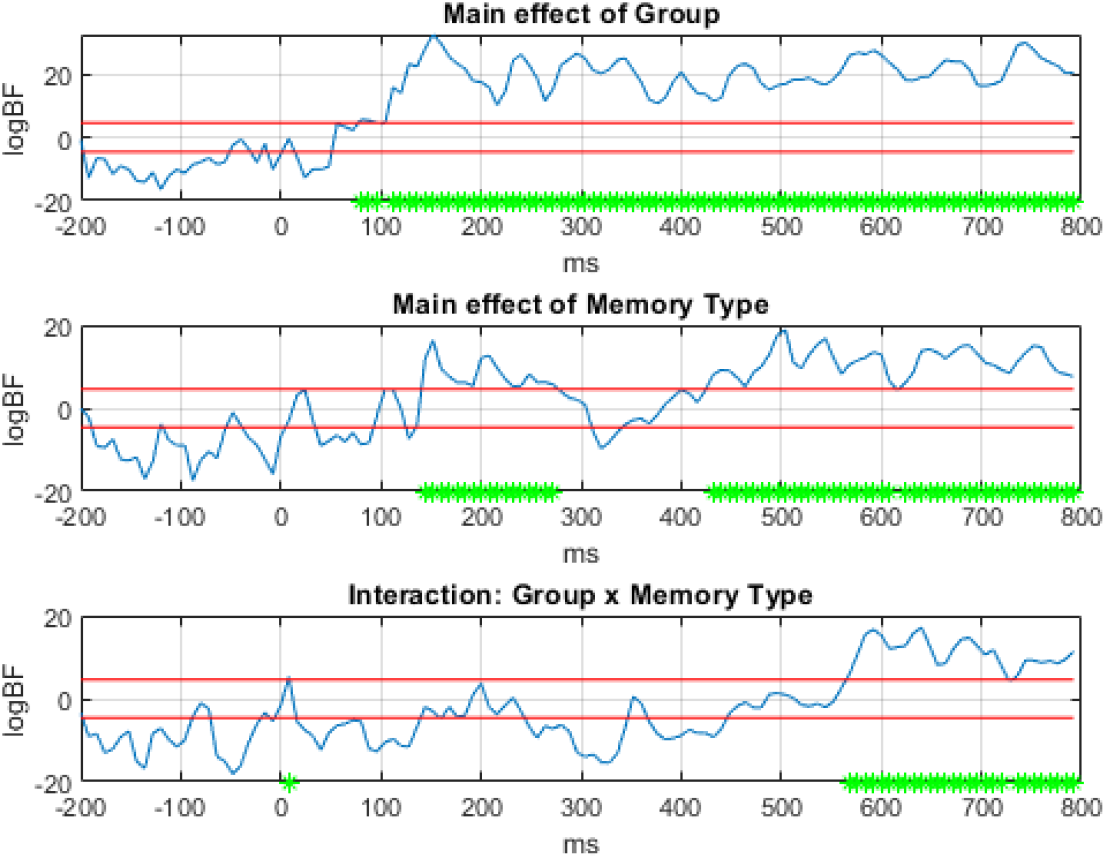
Factorial Analysis of Declarative Memory data - Multivariate Tests. The top to bottom panels plot the Log Bayes Factor in favour of the alternative versus null hypothesis for the main effect of group (old versus young), main effect of memory type (semantic versus episodic) and the interaction. The red lines mean the same as in Fig 5.

For comparison, Fig 12 presents a decoding analysis of the main effects using SVM. MLC and SVM somewhat agree about the main effect of Group but not about the main effect of Memory Type - here, MLC finds an effect but SVM does not. Tables 4 and 5, show that the correlation between MLC Bayes Factors and decoding rates was stronger for the effect of Group than Memory Type (0.86 versus 0.54). This also holds for the correlation between linear and SVM decoders (0.85 versus 0.57).

**Table 4.** Correlations among encoding and decoding scores for main effect of Group. In the top row the correlations are between the log Bayes factor time series from MLC and the *p*_*corr*_ time series from the decoders. All other correlations are among *p*_*corr*_ time series. All correlations are significant at *p <* 0.001.

| Encoder/Decoder | Decoder |  |  |  |
| --- | --- | --- | --- | --- |
|  | kNN | Linear | SVM | PCA-SVM |
| MLC | 0.79 | 0.89 | 0.86 | 0.84 |
| kNN | - | 0.72 | 0.68 | 0.69 |
| Linear | - | - | 0.85 | 0.76 |
| SVM | - | - | - | 0.71 |

**Table 5.** Correlations among encoding and decoding scores for main effect of Memory-Type (episodic versus semantic). In the top row the correlations are between the log Bayes factor time series from MLC and the *p*_*corr*_ time series from the decoders. All other correlations are among *p*_*corr*_ time series. All correlations are significant at *p <* 0.005.

| Encoder/Decoder | Decoder |  |  |  |
| --- | --- | --- | --- | --- |
|  | kNN | Linear | SVM | PCA-SVM |
| MLC | 0.48 | 0.57 | 0.54 | 0.35 |
| kNN | - | 0.25 | 0.44 | 0.46 |
| Linear | - | - | 0.57 | 0.28 |
| SVM | - | - | - | 0.28 |

**Fig 12.**
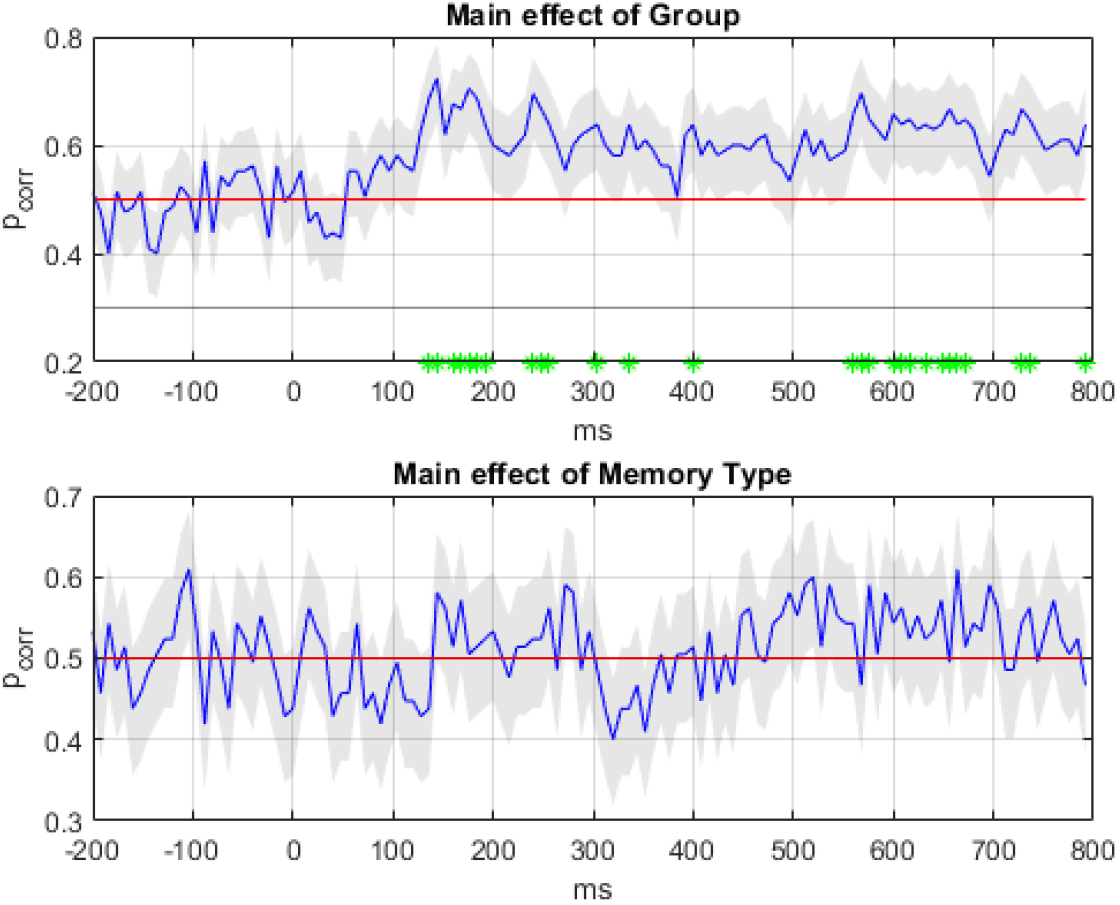
Decoding Analysis of Declarative Memory data. The panels plot the decoding accuracy, *p*_*corr*_, with 90 percent confidence intervals for the main effect of Group (old versus young), and main effect of Memory-Type (semantic versus episodic) using an SVM decoder. The green asterisks denote significant decoding periods (*FDR <* 0.01).

To help understand the MLC results Fig 13 plots Sensor Maps from univariate analyses at the time points corresponding to the strongest multivariate effects. These are chosen at the peak of the Group effect (t=152ms), the peak of the Memory-Type effect (t=504ms) and the peak of the Interaction (t=640ms). The most striking finding is that there are no univariate effects of Memory-Type at t=504ms. Given that SVM decoding also finds no effect, this suggests that the MLC results are false positives.

**Fig 13.**
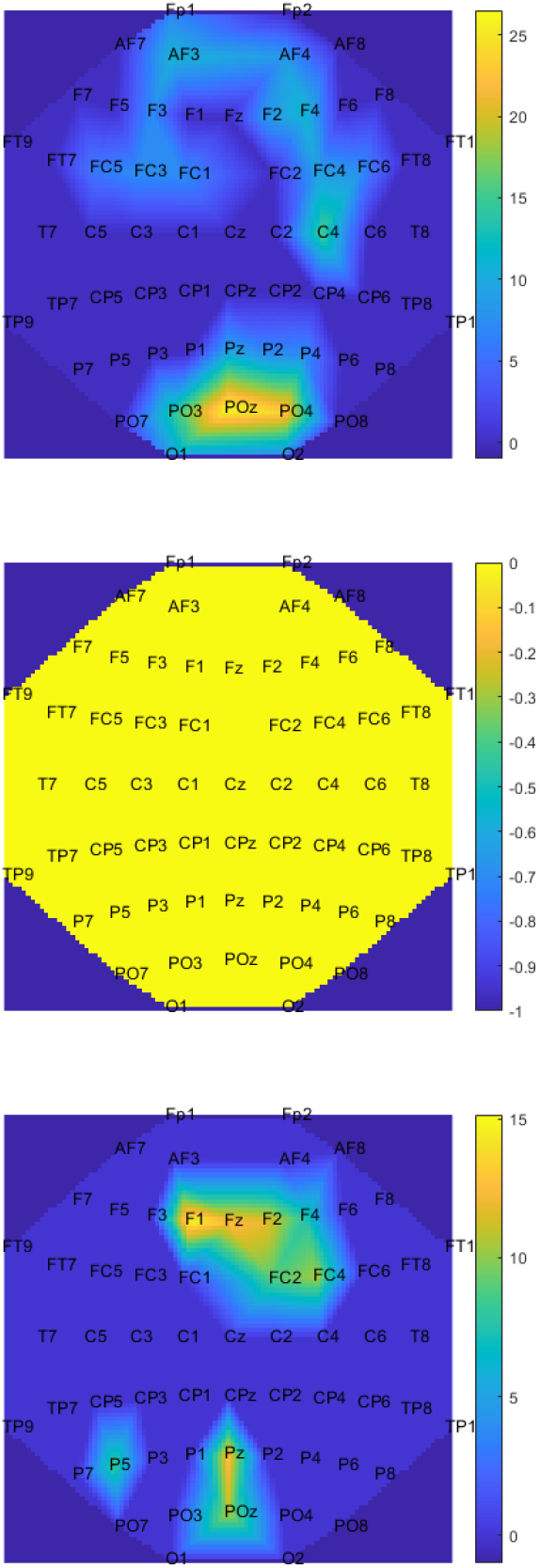
Factorial Analysis of Declarative Memory data - Univariate Tests. The top to bottom rows are for the main effect of Group (old versus young) at 152ms, main effect of Memory Type (episodic versus declarative) at 504ms and the interaction at 640ms. These times correspond to the peaks in Fig 11. The colours indicate the value of the F-statistic from a voxel-wise ANOVA. F-values are set to zero for voxels that do not satisfy an FDR correction over electrodes (*FDR <* 0.05) The yellow image in the middle indicates there are no univariate effects of Memory Type at this time point.

However, this is not the case. In follow-up analyses we calculated what the contribution to the Bayes Factor was from covariance terms alone, as opposed to variance terms. This was implemented by comparing the reported Log Bayes Factors to those obtained after diagonalising Ψ_*n*_ (see “The Nature of Multivariate Effects” in the Methods section). This produced a change of −32.2. The multivariate Memory-Type effects at this time point are therefore due to Voxel Dependencies (VDs) rather than to a Collection of Univariate (CU) effects. We return to this issue in the discussion.

Moreover, we repeated this analysis at time points for which VDs were not driving the inference (ie. where the above difference was zero or negative), and indeed found significant regions in the Sensor Maps (therefore implying CU effects). It therefore appears to be the case that SVMs are not very good at using VDs to drive classification. Because Principal Component Analysis captures voxel dependencies we then had the idea to first project the spatial EEG data onto the first 8 principal components and then run SVM decoding. We then indeed found significant decoding accuracies for Memory Type at similar times as for MLC, as shown in Fig 14.

**Fig 14.**
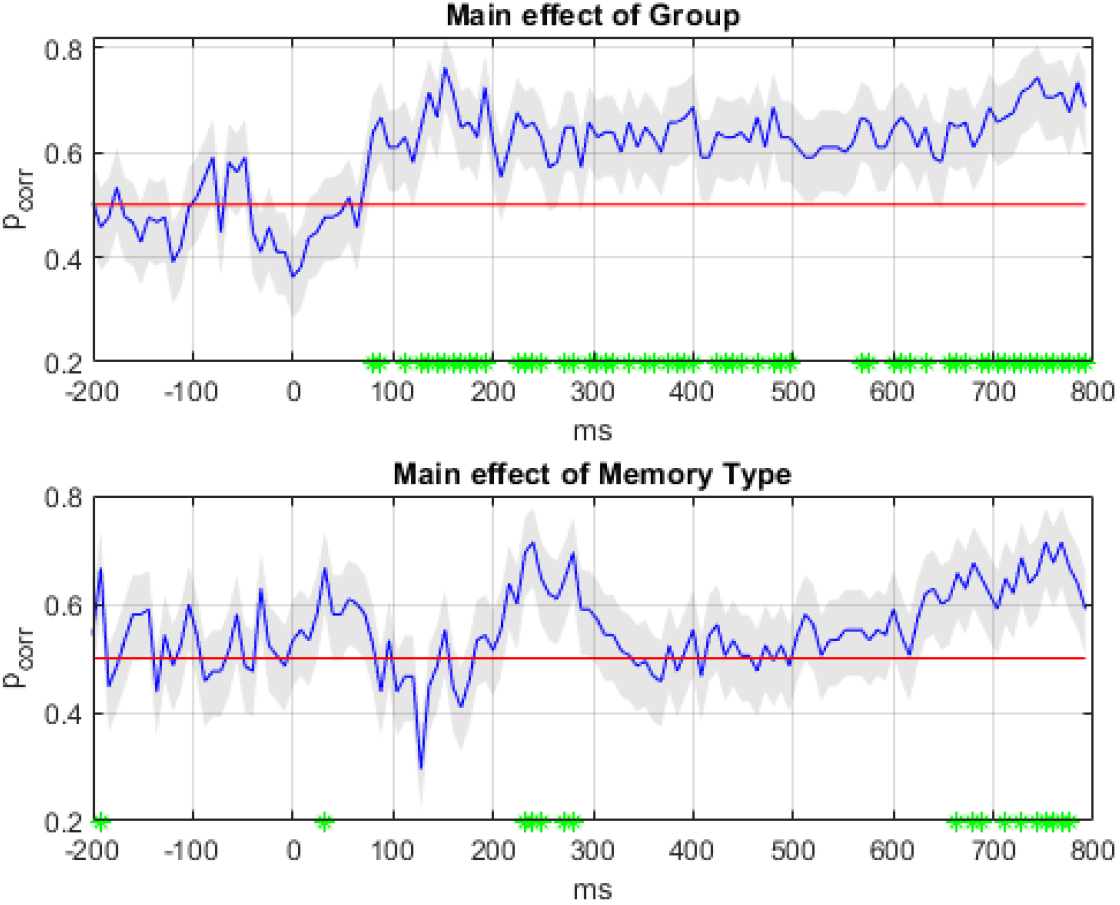
PCA-based Decoding Analysis of Declarative Memory data. The panels plot the decoding accuracy, *p*_*corr*_, with 90 percent confidence intervals for the main effect of Group (old versus young), and main effect of Memory-Type (semantic versus episodic) using an SVM decoder after projecting the data at each time point onto its first 8 principal components. The green asterisks denote significant decoding periods (*FDR <* 0.01).

However, the low correlations between MLC and PCA-SVM shown in Table 5 perhaps suggest that MLC is better able to switch between being driven by CU versus VD effects, depending on the nature of the effects present at each time point.

## Discussion

This paper has shown how a Multivariate Linear model with Conjugate priors (MLC) can be used for the factorial analysis of EEG time series data. Although this model is well known in the field of Multivariate Statistics [33] and Bayesian Econometrics [34] we are unaware of any empirical applications that make use of the associated Bayes factors. We have provided proof of principle of the approach on simulated data and applied it to within-subject analysis of reward learning ERPs and between-subject analysis of declarative memory ERPs.

Computation times for MLC are similar to established decoding methods such as SVM. This indicates that MLC will also be suitable for the analysis of higher-dimensional data such as MEG. The major computational bottleneck for MLC as applied in this paper is in the calibration algorithm which tunes the expected variance of regression coefficients on baseline data. It is possible that alternative methods of setting this parameter can be found. The second bottleneck is computing the determinant of the posterior scale matrix, Ψ_*n*_, that captures the error covariance structure. This is implemented using Singular Value Decomposition (SVD) [50]. For applications of MLC in which the number of independent variables is greater than the number of dependent variables the computational bottleneck will be in inverting and computing the determinant of the regression coefficient covariance matrix, *V*_*n*_.

### Interactions

We envisage that one of the most useful applications of MLC will be to the identification of interactions in multivariate data. We have shown how this is possible using simulated data, and data from ERP studies of reward learning and declarative memory.

We note that there is already a well-established methodology for detecting multivariate interactions in neuroimaging data, namely Dynamic Causal Modelling [51]. Advantages of DCM are that you can make inferences about quantities of neurobiological interest such as changes in connectivity among brain regions. However, DCM does make rather specific assumptions about the functional form of the underlying neurodynamics. Some users may see this as a drawback but others will see this as useful integration of neurobiological knowledge.

Finally, the usefulness of factorial experimental designs, that they efficiently test for the effects of multiple experimental factors and their interactions on an experimental outcome, is very well established across decades of research in multiple fields of enquiry. There is no reason why they should be less useful when that outcome is multivariate. The focus on decoding, however, has perhaps distracted the field from this important point.

### Bayes Factors versus Decoding Accuracy

Our results showed significant correlations between the Bayes factors from MLC and decoding accuracy. These correlations were strong for the within-subject Reward Learning data but less so for the between-subject Declarative Memory data. This may be because decoding at the between-subject level is a more challenging task with typically lower decoding accuracies [5]. These results add to a body of work which finds strong correlations between Bayes Factors and other established measures of model performance. This includes Multilayer Perceptrons [52] and Source Reconstruction models [53] (cross-validated accuracy), Dynamic Causal Models (Akaike’s Information Criterion, Bayesian Information Criterion) [54], and t-tests [55] and ANOVAs [56] (log p-values).

### Within-Subject effects at the Group Level

In future work we plan to run a within-subject analysis on all subjects from the reward learning experiment and make inferences at the group level using Random Effects Bayesian Model Comparison (RFX-BMC) [57]. This will compute, at each time point, the frequency with which main effects or interactions are found in the sample from which the group of subjects are drawn (e.g. 80% of subjects). A plot of the “exceedance probability” will then show at which time points this frequency is significantly higher for the alternative hypothesis than for the null (or vice-versa).

### What drives multivariate effects?

We have reviewed a previous proposal that the power of multivariate versus univariate inference in neuroimaging is that, although effects may be weak at individual voxels, if enough voxels are showing these effects then the resulting multivariate effect can be significant [58]. Pakravan et al [40] refer to this as a Collection of Univariate (CU) effects. We note that if CU effects are local and spatially contiguous then they should also be detectable using (parametric or non-parametric) cluster-level inference. Multivariate inference extends the detection of CU effects to spatially non-contiguous regions as shown for example in the multiple clusters in Figs 9 (bottom) and 13 (top and bottom).

However, it is also possible that multivariate effects are driven by Voxel Dependencies (VDs). The support for this from fMRI is mixed. Pakravan et al. [40] refer to this as coordinated multivoxel coding but find little evidence of it in their analysis of fMRI data from visual processing studies in humans and macaques. But, based on experimental and computational studies of correlations in neural variability, Bejjanki et al [41] make two proposals. The first is that when neurons in a population are selective for the same stimulus then lower noise correlations between them allow increased information to be extracted. This is precisely the relationship that we have derived for MLC in the “The Nature of Multivariate Effects” section (see Methods). The second proposal is that, when neurons in a population are selective for different stimuli, then *increased* correlation among them allows decision boundaries to be more accurately identified. Evidence for this second proposal was provided from decoding analyses of human fMRI data from an attentional control study.

Our results on the between-subject analysis of the Declarative Memory EEG data have shown that MLC is sensitive to both CU and VD effects and that the degree to which they are sensitive to the latter versus former can be quantified by diagonalising the posterior scale matrix that captures covariances, Ψ_*n*_, and reporting the resulting change in log Bayes Factors.

In terms of comparisons with previous univariate analysis [49] of this data (using pre-selected sets of electrodes) the results for the main effect of memory type are largely coherent with the two time periods chosen in [49] and the findings of interactions between memory and age are consistent in that effects were found in the 500-800ms window but not earlier (see Fig 11).

### Accepting the Null

Bayesian approaches to model inference allow for the null hypothesis to be accepted and have previously been developed for the factorial analysis of univariate data [56]. With the multivariate reward learning data, the textbook expectation is that there will be neural correlates of Reward Prediction Errors (RPEs) per se [59]. This mandates that there will be an interaction between the sign and magnitude of the RPE because, for positively signed RPEs, increasing RPE will increase value, but for negatively signed RPEs, increasing RPE will decrease value. Acceptance of the interaction null hypothesis when the main effects can be accepted (see Fig 8) is therefore an indication that some other valuation process is active [60, 61], for example, a valence encoder (main effect of sign), a salience encoder (main effect of magnitude) or both effects together (presumably with distinct neural sources).

### Accuracy of Differences

The standard way to test for interactions using multivariate decoding is what we have referred to as a “difference of accuracies” test. We have given an examples of its application in various EEG/MEG contexts [19–22]. We emphasise that the statistically significant findings from these studies are completely valid. However, as we have argued earlier, negative findings are uninterpretable as there can be interactions without differences in accuracy.

We now propose that those wishing to pursue tests for interactions using decoding methods use an “accuracy of differences” test which we now define. This is most naturally implemented at the between-subjects level and is based on difference images. Given a 2-by-2 design, for each subject, one first creates a difference image *d*_1_ which is the difference between responses at the first level of the second factor (e.g. *d*_1_ = *Y A* − *Y B* for our simulated example) and *d*_2_ at the second level of the second factor (e.g. *d*_2_ = *OA* − *OB*). One then tests whether the accuracy for decoding *d*_1_ versus *d*_2_ is significantly above chance. For a *k*_1_-by-*k*_2_ design the relevant difference images and accuracy tests can be informed by the contrast matrices used to define interactions in linear models [42]. However, with more than 2 levels of a factor, there will be multiple accuracy tests and only one need be significant for there to be an interaction. This adds a degree of complexity. Moreover, for inference at the within-subjects level, it is not immediately clear how to pair the trials to create the difference images (at the between-subject level there is only a single image per condition per subject so this ambiguity does not arise). Possibilities include pairing trials closest in experimental time, or using bootstrap resampling to make inferences over multiple pairings. However, we leave the development of “accuracy of differences” tests for others to pursue.

## Materials and methods

This section presents a Bayesian Multivariate Linear model with Conjugate priors which we refer to as MLC. We review the underlying statistical assumptions, inference procedure, computation of Bayes factors, properties of Bayes Factors, and describe how MLC can be calibrated to run MANOVAs for EEG time series data. We also describe the decoding methods used in the empirical work and the data sets that the analysis methods are applied to.

### Multivariate Linear Model with Conjugate Priors

Although this model is well known in the field of Multivariate Statistics [33] and Bayesian Econometrics [34] we are unaware of any empirical applications that make use of the associated Bayes factors. The mathematical basis of our implementation is described in [35] and our notation makes use of the Matrix-Normal and Matrix-T distributions defined in Gupta and Nagar [36]. For completeness, the definitions of these distributions and their properties are also described in S1 Appendix.

#### Likelihood

The Multivariate Linear Model is given by

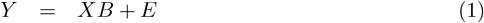

where *Y* is a [*N* × *M*] matrix of dependent variables, *X* is an [*N* × *K*] matrix of independent variables, *B* is a [*K* × *M*] matrix of regression coefficients and *E* is an [*N* × *M*] matrix of Gaussian errors. Each multivariate Gaussian noise sample (*e*_*n*_, row of E) has zero mean and [*M* × *M*] covariance matrix Σ. We can write this as

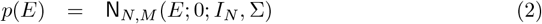

where N_*N,M*_ is a Matrix-Normal density, and *I*_*N*_ indicates that the *N* trials are assumed independent and identically distributed. The Matrix-Normal density has three parameters; a matrix of mean values, the covariance over rows and the covariance over columns. One can write this distribution as a product of multivariate Normal densities (that is, densities over vectors) but the matrix form provides a compact notation for the likelihood, prior and posterior densities (see S1 Appendix for further details). The likelihood is therefore

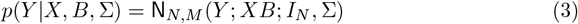

with mean *XB*.

#### Priors

We assume a Normal-Inverse Wishart prior

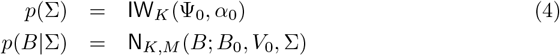

where *B*_0_ is the prior mean, *V*_0_ is the [*K* × *K*] prior covariance matrix over rows of *B*, and Σ is the prior covariance matrix over the columns of *B*. The marginal prior over the regression coefficients is therefore

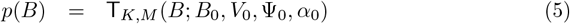

where T_*K,M*_ is a Matrix-T distribution. These are conjugate priors, meaning that the priors and posteriors have the same mathematical form. The Matrix-T distribution has four parameters; a matrix of mean values, covariance over rows, scale matrix over columns, and degrees of freedom (see S1 Appendix for further details).

#### Posteriors

The posteriors are

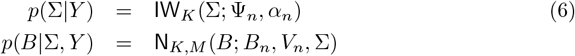

The marginal posterior over regression coefficients is

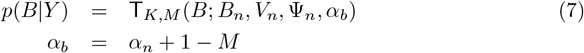

We have

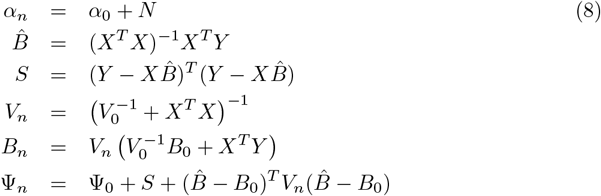

with posterior mean *B*_*n*_, posterior row covariance *V*_*n*_, and posterior column scale Ψ_*n*_. Many of the above quantities are likely familiar to the reader with 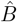 equal to the Maximum Likelihood estimate of the regression coefficient matrix, and *S* corresponding to the matrix of Sum-Squared Errors.

#### Bayes Factors

The Bayes factor is defined as the ratio of model evidences under the alternative versus null models

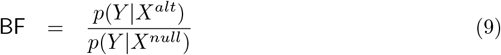

See below section on Factorial Designs for how we set up the design matrices. The log Bayes Factor in favour of the alternative versus null model (see S1 Appendix) is then

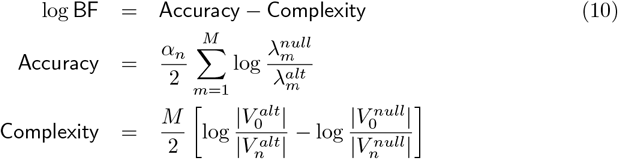

where 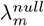 and 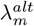 are the eigenvalues of 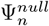 and 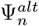 respectively and correspond to error variances in the *m*th eigen-direction. The notation |*V*| denotes the determinant of matrix *V*.

The accuracy term is larger if the error volume (given by the product of eigenvalues - see S1 Appendix) is smaller under the alternative model. This is similar to (but not the same as) classical measures employed in MANOVA such as Pillai’s statistic which measures the ratio of explained to total variance [23]. The complexity term is equal to the information gained about the regression coefficients. This derives from the fact that the entropy of a Gaussian variable with covariance *V* is equal to log |*V* | (plus constants) [37]. Thus each term corresponds to entropy lost (or information gained) under the respective models. Importantly, we emphasise that computation of the Bayes factor is exact and analytic. Approximate inference procedures such as Variational Bayes [32] or Markov Chain Monte Carlo [38] are not required.

We can accept the alternative hypothesis if the Bayes Factor exceeds a specified threshold. Here can we make use of the fact that a posterior probability of *p*_*T*_ corresponds to a False Discovery Rate (FDR) of 1 − *p*_*T*_ [39]. Given that the posterior probability is, in turn, a sigmoid function of the log Bayes Factor, we can enforce FDRs of 1% with log Bayes Factor thresholds of 4.6. To accept the Null requires a log Bayes Factor threshold of −4.6. These are the thresholds used for the empirical work in this paper.

### The Nature of Multivariate Effects

This section describes how we can understand what signal differences are driving inferences about multivariate effects. We first consider a simplified example in which the covariance of signals about their mean, as captured by the posterior scale matrix, Ψ_*n*_, has a simplified form. Specifically, if Ψ_*n*_ is equal to a scalar term *σ*_*e*_ multiplied by a Toeplitz matrix (in which first neighbours have correlation *r*_*e*_, second neighbours 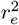, third neighbours 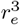 etc.) that reflects a spatial correlation of *r*_*e*_, then entering this into equation 10 produces two separate contributions to model `Accuracy`. First, the scalar term contributes

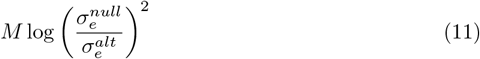

so is in favour of the alternative if 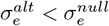. This term corresponds to what has been described as a Collection of Univariate (CU) effects (see [40] and Discussion). Second, the correlation term, depicted in Fig 15, shows that the contribution is in favour of the alternative if the errors are less correlated than under the null 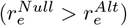. This is analogous to observations from experimental and computational studies [41] which show that, if neurons in a population are selective for the same stimulus, then lower noise correlations between them allow increased information to be extracted. It also corresponds to what has been described as Voxel Dependency (VD) effects (see [40] and Discussion).

**Fig 15.**
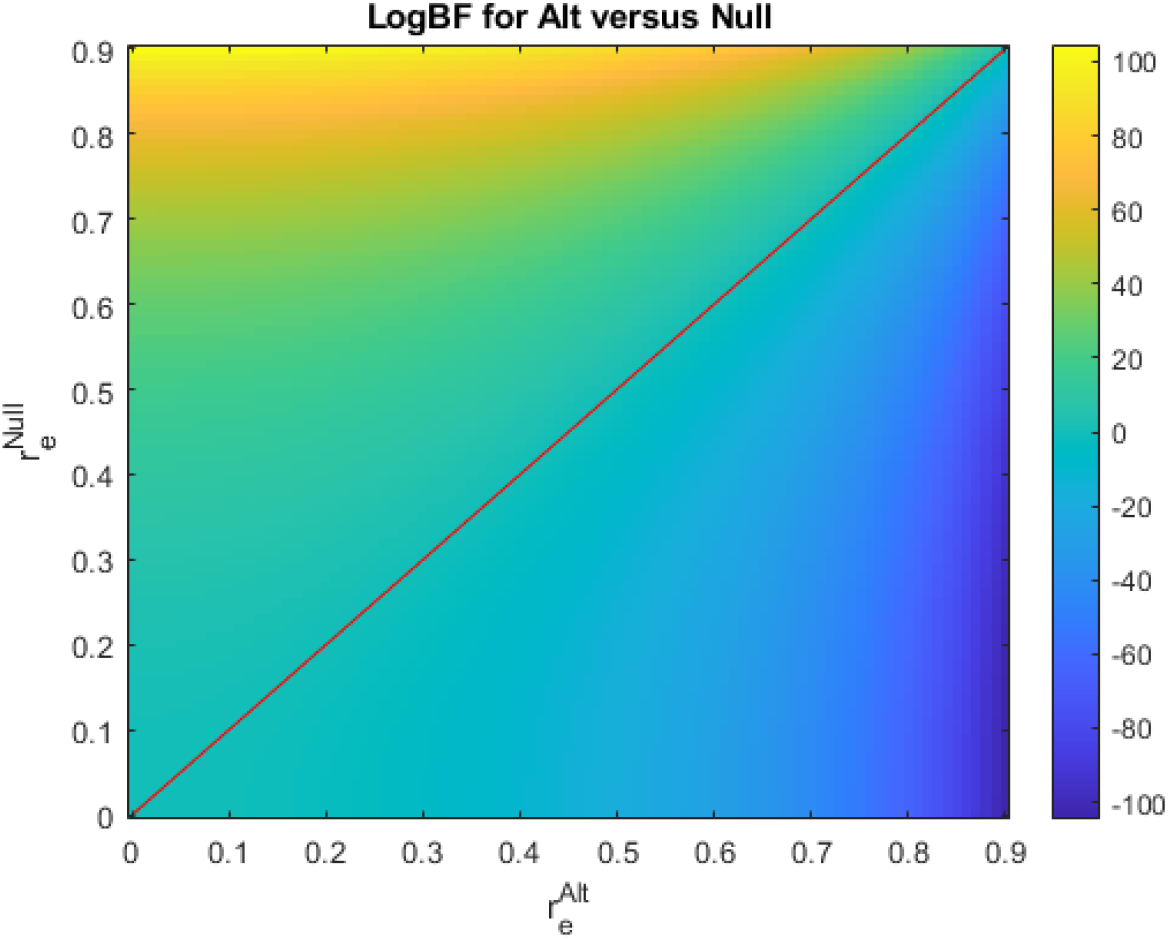
Contribution to Log Bayes Factor from error correlations over electrode space. The contribution is in favour of the alternative if the errors are less correlated than under the null (if 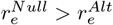 which is the area above the red line).

Given empirical data we propose a straightforward method for quantifying the degree to which inferences are driven by CU versus VD effects which is simply to diagonalise the estimated Ψ_*n*_ matrix and compute the change in Bayes factor. The reduction indicates the degree to which inference is driven by VD effects and we shall see an example of this in the Results section.

#### Calibration

In this paper we use the following priors

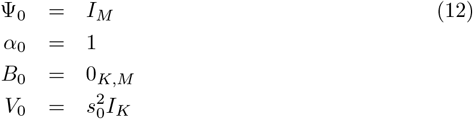

For a Wishart distribution to have well-defined moments the degrees of freedom need to be *α*_0_ *> M* + 1. For the empirical results in this paper, however, we used *α*_0_ = 1 and found this to work well in practice.

The only parameter we need to specify is *s*_0_ which sets the expected magnitude of regression coefficients. We set this as follows. We apply MLC to time series data from EEG experiments in which it is known that there are no effects in the so-called ‘baseline period’, which is typically defined as the one or two-hundred milliseconds before the event of interest occurs (around which the trials have been epoched). We then set *s*_0_ so that the largest log Bayes factor is less than zero, but as close to it as possible. This is implemented using an iterative calibration algorithm which starts with *s*_0_ = 1 and reduces it in steps of *a* × *s*_0_ with step size *a* initially set to 0.5. If we overshoot, then we halve the step size and continue the search from the last valid value. This continues for a maximum of 16 iterations.

#### Factorial Designs

Given a [*k*_1_ × *k*_2_] factorial design there are *k* = 1..*K* experimental conditions with *K* = *k*_1_*k*_2_. If *N*_*k*_ is the number of samples in the *k*th condition then the factorial design matrix *X*_*f*_ has a column for each of the conditions and is a block-diagonal matrix where each block is equal to 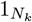 (a column vector of *N*_*k*_ ones).

Bayesian tests for main effects and interactions can then be implemented in one of two ways. First, by fitting the data, *Y*, using MLC with design matrix *X*_*f*_. One then specifies the relevant contrast matrix [42], *C*_*j*_, that implements the *j*th test e.g. *C*_*j*_ = [1, 1, −1, −1] to test for the main effect of factor 1 in a 2-by-2 design. One could then make an inference using a “Savage-Dickey” or “Posthoc” approximation to the model evidence [43] based on the relevant posterior T-distribution. This approach fits a single model and uses a different contrast to test for each experimental effect.

However, as we have an analytic expression for the Bayes factor we use a second approach which fits a separate alternative model for each effect to be tested. We first fit a null model with design matrix *X*_0_ = 1_*N*_ (i.e. constant over conditions) where *N* is the total number of ERPs. The design matrix to test for the *j*th effect is

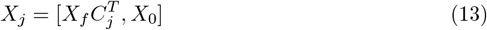

This has the null model in the last column, and preceding columns contain the relevant patterns of differences among conditions (see Fig 4 for an example). The log Bayes factor in favour of an alternative hypothesis is then given by Eq 10. In the above, the null hypothesis is that the effect is zero at all electrodes. To test for main effects and interactions in two-way ANOVAs we fit four models; a null model, a model for each of the main effects and a model for the interaction.

### Computational Bottleneck

We emphasise that estimation of model parameters is analytic and implemented using Eq. 8 only. This results in a fast algorithm because iterative optimisation is not needed. Additionally, although matrix inversions are required they are only over [*K* × *K*] matrices where *K* is the number of independent variables. Computation of the log Bayes Factor requires log determinants for the [*K* × *K*] matrix *V*, and [*M* × *M*] matrix Ψ where *M* is the number of output variables. In this paper *M* is the number of electrodes and is much larger than the the number of variables in the design matrix. But applications with *K > M* will have different scaling properties.

### Decoding Methods

The empirical work in this paper compares MLC with decoding approaches based on Support Vector Machines (SVMs) [44], Linear Decoders and k-Nearest Neighbour [45]. We used the implementations in Matlab’s Statistics toolbox with default options, specifically the functions `fitcsvm.m` which is optimised for low-dimensional data and `fitclinear.m` which is optimised for higher dimensional data (for *M* > 2 we use the function `fitcecoc.m`). For the nearest neighbour classifier we used `fitcknn.m` with *k* = 5. For reasons explained in the Results section we also ran decoding analyses by first projecting the data at each time point onto its first 8 principal components, and then running SVM. This overall approach is referred to as PCA-SVM.

Classification accuracy was estimated using ten-fold cross-validation and confidence intervals were derived using the method of Brodersen et al. [46] which uses the number of correct and incorrect classifications over all folds to define parameters of a beta distribution. The 10th and 90th percentiles of this distribution can then be read off from the associated cumulative density function.

A p-value at each time point was computed based on the classification accuracy and its binomial distribution. Correction for multiple comparisions across time was implemented using the Benjamini-Hochberg procedure [47] for controlling the False Discovery Rate at *FDR* < 0.01 (see green asterisks in decoding Results figures).

### Simulated Data

We created synthetic data from a simulated EEG experiment conforming to a 2-by-2 factorial design with factors of “Group” (young, Y, versus old, O) and “Drug” (A versus B). We first created a data set with a main effect of Drug, no effect of Group, and no interaction. For each group, data was created by adding differences in means (between drug conditions) at 25 per cent of electrodes (same electrodes for all conditions). This difference, *d*_*t*_, had a Gaussian-like temporal profile

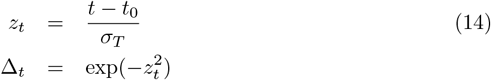

that peaks at *t*_0_ = 267ms with a width of *σ*_*T*_ = 40ms. These differences were the same for both groups (so no effect of Group). The spatial correlation between electrodes *i* and *j* was set to

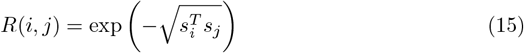

where *s*_*i*_ is the 2D location of electrode *i* on the scalp surface (in normalised units). Gaussian noise was added with mean 0 and covariance 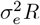 such that the ratio of signal standard deviation to noise standard deviation *σ*_*e*_ was 25 per cent at peak time *t*_0_.

We therefore expect the Drug conditions to be discriminable at approximately between plus and minus 1.6 *σ*_*T*_, which works out to be between 200 and 333ms. We created ERPs for *N*_*c*_ = 26 simulated participants in each group. Images of the mean activity in each condition are shown in Fig 2. We then created a new data set, S2, in which the sign of the difference in Drug conditions changed as a function of Group, as shown in Fig 3. This created an interaction between Group and Drug but removed the main effect of Drug.

The effects shown in Fig 2 and 3 were designed so that the correspondence with the univariate effects described in the introduction is especially clear; these simulations have essentially the same univariate effect replicated at multiple voxels. We refer to these as a homogeneous effects. We also created a data set, S3, containing heterogeneous effects, for example, positive differences at one electrode, negative at another (these were created by flipping the sign of Δ_*t*_ at selected electrodes). Additionally, as with empirical data, we created different effects at different time points by selecting different values of *t*_0_. The mean trajectories from this data set are shown in Fig 16.

**Fig 16.**
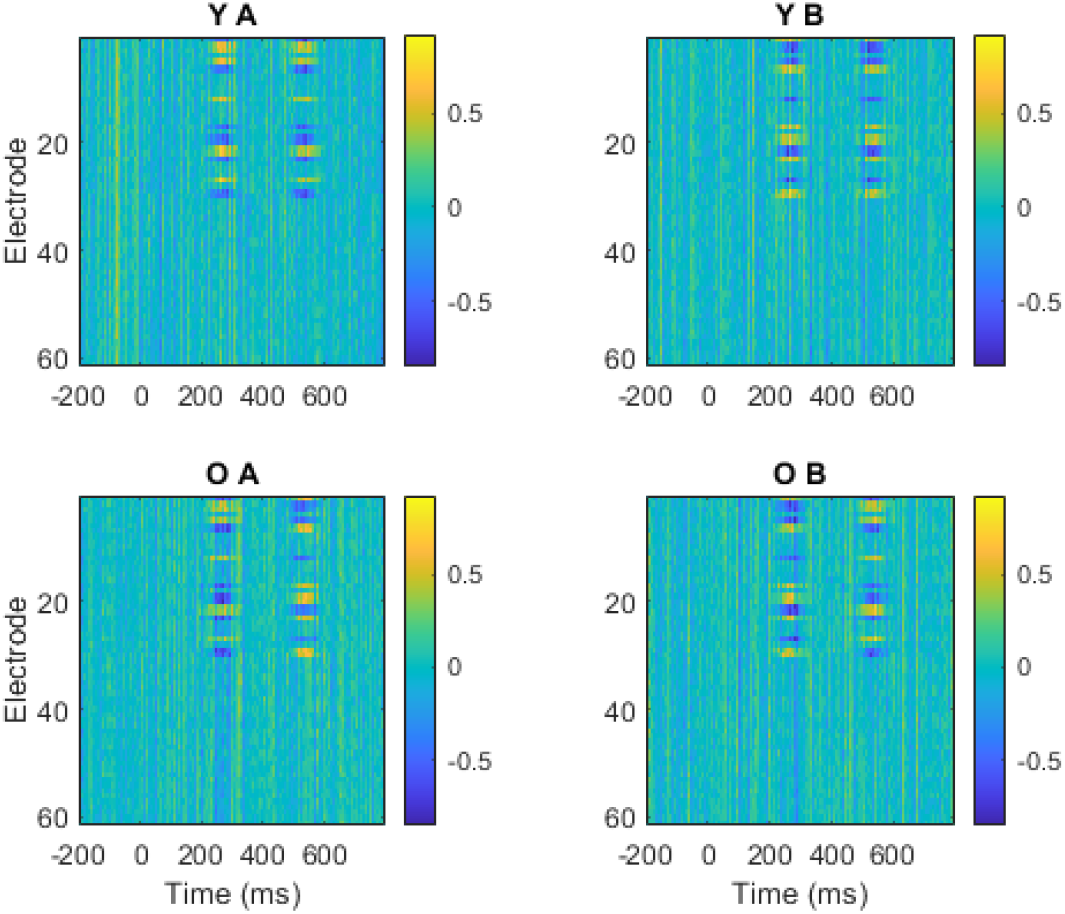
Simulated Data S3. This data set contains heterogeneous responses over electrodes corresponding to a main effect of Drug at 267ms (the patterns are different for drugs A and B and this difference is the same for Y and O groups) and an interaction between Group and Drug at 533ms (the patterns are different for drugs A and B and this difference is different for Y and O groups).

### Reward Learning

The reward learning study [48] was approved by the ethics committee of the Faculty of Health and Human Sciences at the University of Plymouth. The current data set is from the N=42 experiment in which Reward Prediction Errors (RPEs) were manipulated by varying outcome likelihood. Here we use data from a single subject only as we wish to demonstrate how MLC can be used for a within-subject analysis.

Participants took part in a probabilistic reinforcement learning task. On each trial, they selected one of two keys and were then given reward feedback. They were informed that one key gave on average slightly better feedback (by the terms of the block, see below) than the other and that they should pay attention to the feedback they received so they might learn which key to prefer for the block’s duration. Trials were presented in blocks of sixty, and the participant told that at the start of each block the good key would be randomly re-selected, requiring a new learning episode. In fact, feedback was pseudo-randomly predetermined, and the participant could not influence the outcome by key selection. Feedback on a trial was binary: either a one or a six. At the start of each block, a guideline probability for sixes was given, which could take the value 25% (low), 50% (medium) or 75% (high). A corresponding target number of sixes for the block was also given: 15, 30 and 45 respectively. On “gain” blocks, participants attempted to reach or exceed the target number of sixes by the end of the block since they received 2 pounds for doing so and nothing if they did not. In these blocks sixes were thus positive Reward Prediction Errors (RPEs) and ones were negative RPEs. On “loss” blocks, participants attempted to avoid reaching the target number of sixes since they lost 2 pounds for doing so and lost nothing if they did not. Once again, in reality neither key was set to have a higher chance of returning sixes, with feedback pseudo-randomly predetermined.

Stimulus magnitude, RPE sign and domain (gain versus loss) were fully counterbalanced. Fifteen gain domain blocks were run followed by fifteen loss domain blocks (sixty trials per block), with this order reversed for half the participants. Within each domain, five sub-blocks were presented for each of the three baseline probabilities for sixes. The order of these sub-blocks was counterbalanced.

EEG data were collected from 61 Ag/AgCl active electrodes (actiCAP, Brain Products, Gilching, Germany) mounted on an elastic cap and arranged in a standard International 1020 montage. EEGs were time-locked to 100 ms before the onset of the feedback to 700 ms afterward, and then were baseline-corrected. Reward Prediction Errors were derived from a Rescorla-Wagner learning model with a learning rate set to the same standard value over subjects. See [48] for further details of preprocessing and behavioural modelling.

For the empirical work in this paper, we analysed data from a single subject (subject “38”) with *N* = 411 trials. This was reduced by a factor of 4 from the atypically large original number of trials in order to speed model fitting. We dichotomised RPEs into “small” versus “large” levels via a median split. The other factor is RPE sign with levels “negative” and “positive”. We therefore analyse the data as a 2-by-2 factorial design. Fig 17 plots the grand mean ERP trajectories for the corresponding experimental conditions.

**Fig 17.**
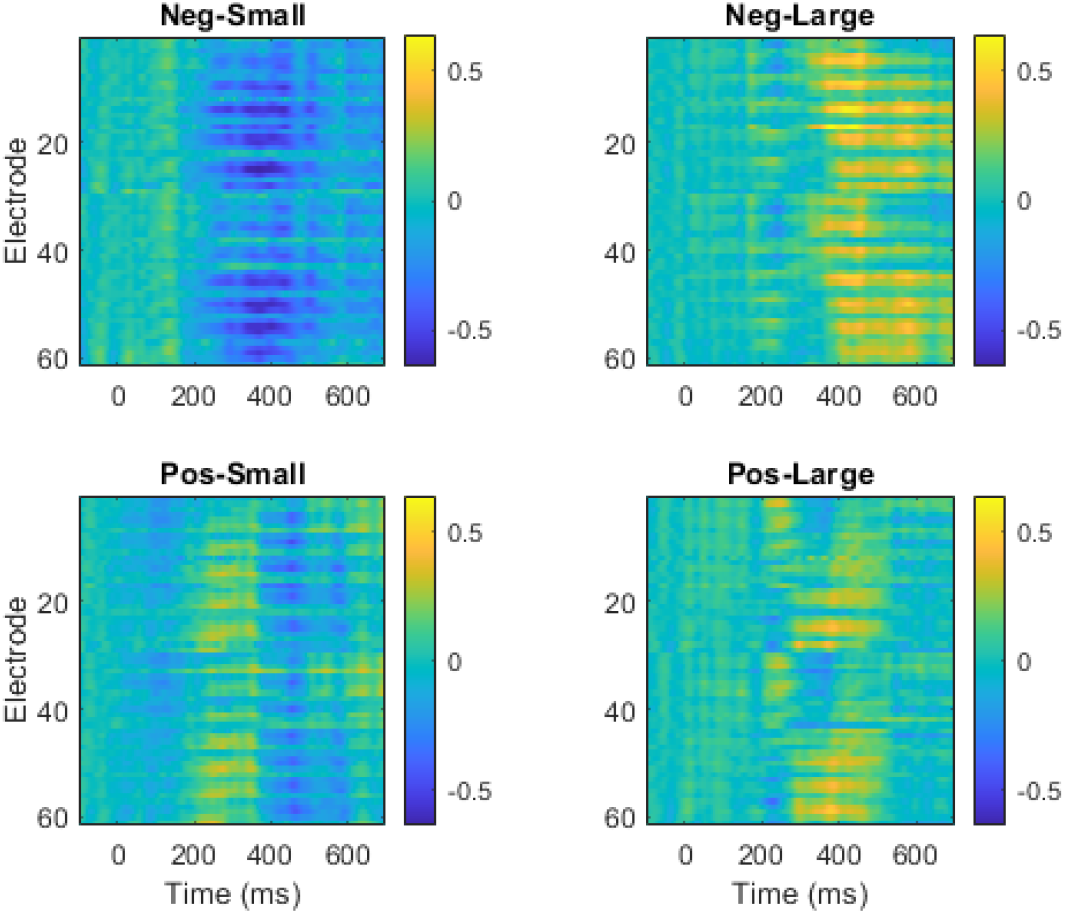
Reward Learning data. Images of mean ERP trajectories in four experimental conditions for a single subject with the top row for Negative RPEs, bottom row for Positive RPEs, left column for small RPEs and right column for large RPEs.

### Declarative Memory

The Declarative Memory (DM) study was approved by the ethics committee of the Psychology School at the University of East Anglia and comprises an ERP data set recorded from 26 young and 26 older adults whilst they categorised personality traits for their relevance to certain groups of people (semantic memory) [49]. For example the trait “fearless” may describe a soldier but not a librarian. Data was also acquired for an episodic memory condition based on earlier study of word lists. We use this data set to show how MLC can be used for a factorial analysis of ERP data at the between-subject level.

EEG was acquired from a 63-channel BrainProducts active electrode system embedded in a nylon cap. An average reference was used and EEG data were segmented into epochs of 1s from −200ms to 800ms after the onset of word stimuli, and the ERP is sampled at 500Hz. We baseline corrected this data within-subject so that the average activity in the baseline period (−200 to 0ms) was zero. Data were multiplied by a scalar value so that the average standard deviation over channels and conditions was unity. Fig 18 plots the grand mean ERP trajectories for the four experimental conditions. In Tanguay et al. [49], ERP analysis focussed on two time periods of interest; the “N400”: 250 to 500ms, and the Late Positive Component (LPC): 500 to 800ms.

**Fig 18.**
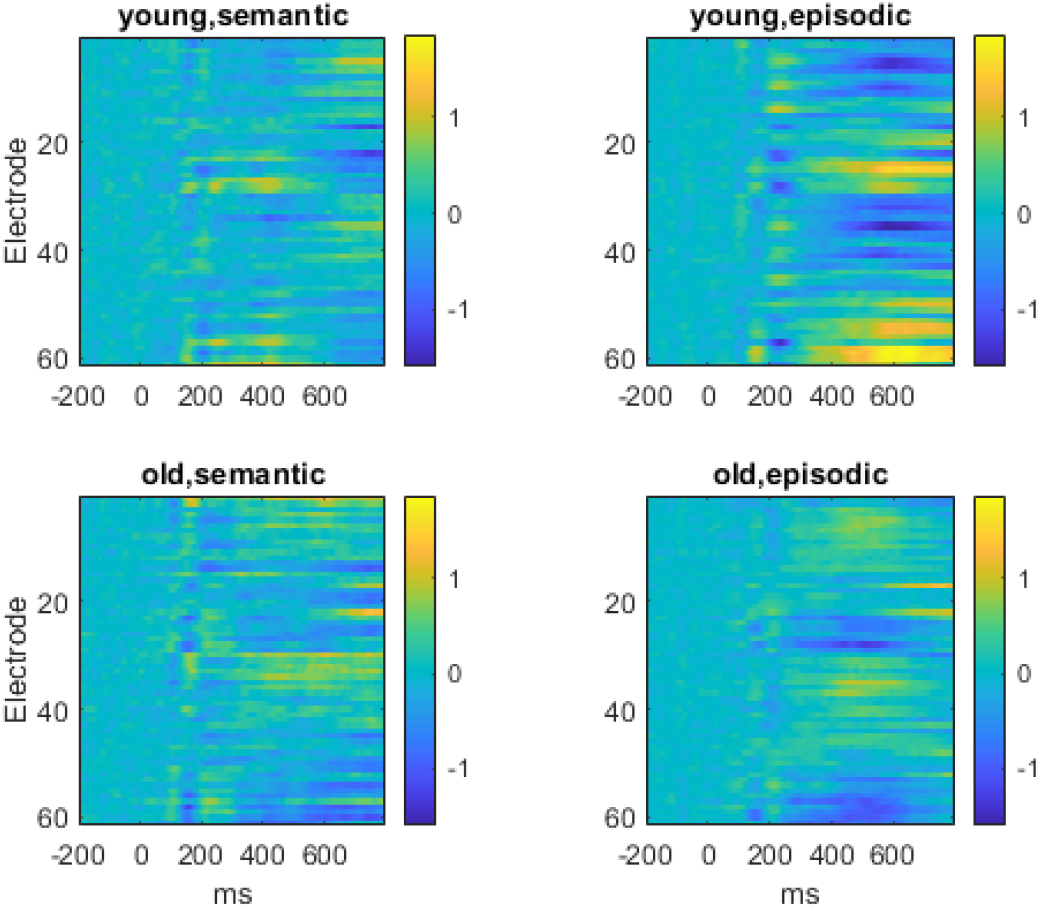
Declarative Memory Data. Images of mean ERP trajectories in four experimental conditions.

## Ethics Statement

This paper contains analysis of secondary data from two previously published studies. The Reward Learning study [48] was approved by the ethics committee of the Faculty of Health and Human Sciences at the University of Plymouth in 2013. Testing was conducted between 1st December 2013 and 6th February 2014 and consent was provided in writing. The Declarative Memory (DM) study [49] was approved by the ethics committee of the Psychology School at the University of East Anglia (REF: 2019-0174-001552). Written consent was obtained from all participants and testing was conducted between 26 February 2020 and 1st October 2022.

## Supporting information

**S1 Appendix. Mathematical Background**. Provides definitions and properties of Matrix-Variate densities, Bayes Factors and Predictive Density for the Multivariate Linear Model with Conjugate Priors (MLC).

## Acknowledgments

We would like to thank Ann-Kathrin Johnen from Birmingham City University for retrieving and reformatting the Declarative Memory dataset, and Nicholas Menghi from the Max Planck Institute for Human Cognitive and Brain Sciences in Leipzig for discussing decoding-based interaction tests.

## 1 Distributions over Matrices

The following definitions and properties are taken from [1].

### 1.1 Normal distributions

The Matrix-Normal distribution over the [*d*_1_ × *d*_2_] matrix *X* is

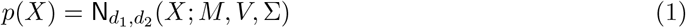

with *M* being the mean, *V* being the [*d*_1_ × *d*_1_] covariance over rows, and Σ being the [*d*_2_ × *d*_2_] covariance over columns. It is equivalent to the multivariate Normal

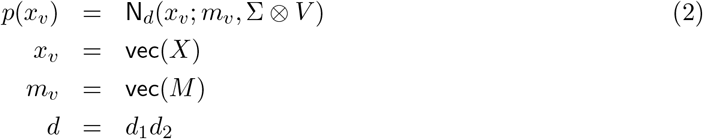

where ⊗ denotes the Kronecker product, x = vec(X) vectorises *X* column by column, and N_*d*_(*x*; *m, C*) is a multivariate Normal distribution over *d*-dimensional random vector *x* with mean *m* and co-variance *C*.

## 1.2 Wishart distribution

The Inverse-Wishart Distribution over the [*p* × *p*] matrix Σ is given by

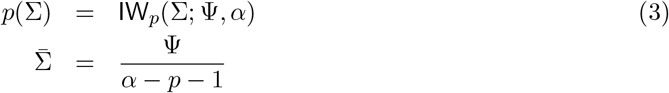

with mean 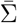 if *α > p* + 1. Its equivalent to stating the Σ^−1^ follows a Wishart distribution. We have

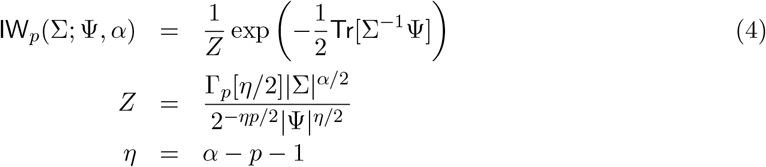

### 1.3 Matrix T-distribution

The Matrix-T distribution over the [*d*1 × *d*2] matrix *X* is

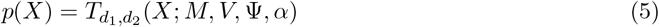

with mean *M* (if *α >* 1 otherwise undefined), [*d*_1_ × *d*_1_] scale matrix *V*, [*d*_2_ × *d*_2_] scale matrix Ψ, and degrees of freedom *α*. We have

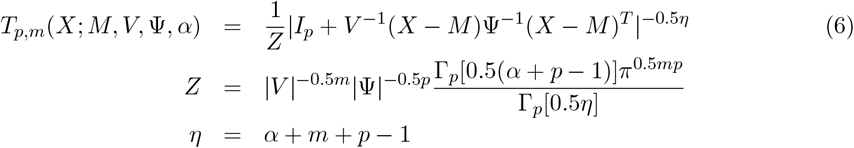

Note that evaluating this density involves computing the inverses *V* ^−1^ and Ψ^−1^.

If *C* = *AXB* where *A* is a [*c*_1_ × *d*_1_] row contrast matrix and *B* is a [*d*_2_ × *c*_2_] column contrast matrix then, according to Theorem 4.3.8 in Gupta and Nagar, we have

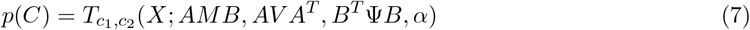

According to Theorem 4.3.3 we have

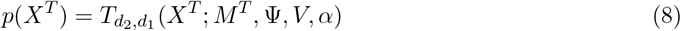

Another property (Theorem 4.3.1) is that

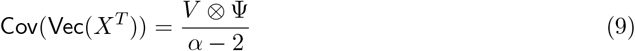

for *α >* 2. So we should therefore have

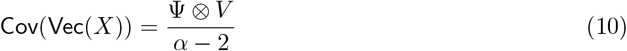

This then matches up with the equivalent formula for the Matrix Normal.

### 1.4 Multivariate T-distribution

The multivariate T distribution T_*d*_(*x*; *m*, Σ, *α*) over *d*-dimensional random vector *x* has mean *m*, scale matrix Σ, degrees of freedom *α*, and covariance

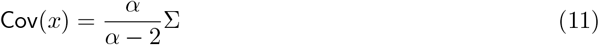

Given [*d* × *r*] contrast matrix *C* we have

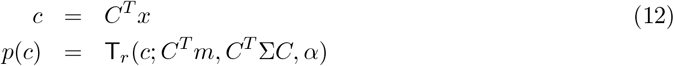

## 2 Model Evidence and Bayes Factors

The model evidence for the Multivariate Linear Model with Conjugate Priors (see main text) is

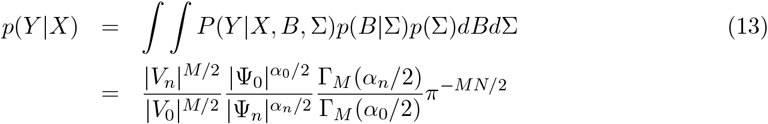

where |*V*_*n*_| is the determinant of *V*_*n*_ and Γ_*M*_ is the generalised gamma function [1]. A derivation is provided in section 3.4 of Ramirez-Hassan’s book [2] which is available digitally at https://bookdown.org/aramir21/IntroductionBayesianEconometricsGuidedTour/.

The Bayes factor is therefore

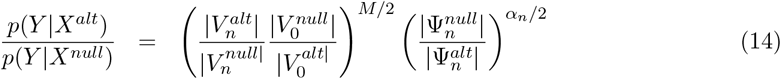

where we have assumed that 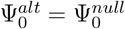 (this is the assumption for our application to EEG data but can be relaxed for other applications). One way to understand the second term is to write it as

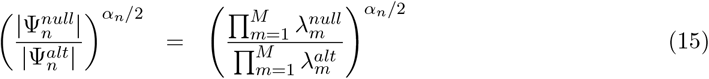

where 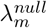 and 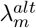 are the eigenvalues of 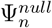 and 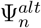 respectively. These correspond to error variances in the *m*th eigen-direction. The product of these eigenvalues corresponds to the error volume in *M*-dimensional space (see Fig 2.7 in [3] for an illustration). The alternative model is to be preferred if it has a smaller error volume. The log Bayes Factor in favour of the alternative versus null model is then

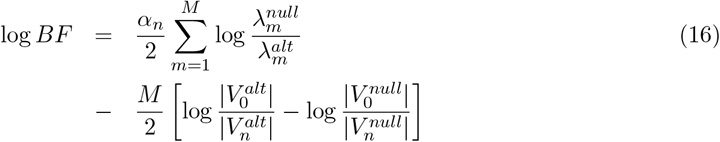

## 3 Predictive Density

The following property is not made use of in the current paper but may be useful in future applications. Given *N*_0_ new observations *Y*_0_ the predictive density is

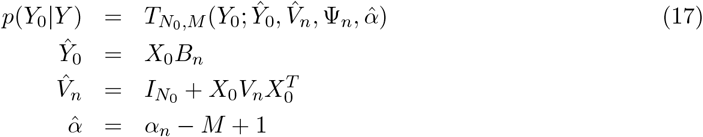

and Ψ_*n*_ is defined in the main text under “Posterior Density”. The predictive density could be used as part of a Bayesian inference approach to infer diagnostic labels from clinical imaging data where *Y* is the training data, *Y*_0_ are the new data to label, *p*(*C*) is a prior distribution over labels, and the posterior distribution over labels is

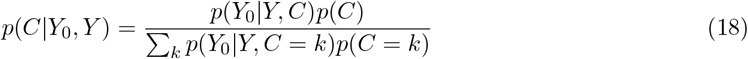

and *P* (*Y*_0_|*Y, C*) takes the form of Eq 17 with parameters (*α*_*n*_, *B*_*n*_, *V*_*n*_, Ψ_*n*_) specific to label *C*.

